# Immunity priming uncouples the growth-defense tradeoff in tomato

**DOI:** 10.1101/2022.07.24.501304

**Authors:** Meirav Leibman-Markus, Anat Schneider, Rupali Gupta, Iftah Marash, Dalia Rav-David, Mira Carmeli-Weissberg, Yigal Elad, Maya Bar

## Abstract

Plants have developed an array of mechanisms to protect themselves against pathogen invasion. The deployment of defense mechanisms is imperative for plant survival, but can come at the expense of plant growth, leading to the “growth- defense trade-off” phenomenon. Following pathogen exposure, plants can develop resistance to further attack. This is known as induced resistance, or priming. Here, we investigated the growth-defense trade-off, examining how defense priming via Systemic Acquired Resistance (SAR), or Induced Systemic Resistance (ISR), affects tomato development and growth. We found that defense priming can promote, rather than inhibit, plant development, and that defense priming and growth tradeoffs can be uncoupled. Cytokinin response was activated during induced resistance, and found to be required for the observed growth and disease resistance resulting from ISR activation. ISR was found to have a stronger effect on plant development than SAR. Our results suggest that growth promotion and induced resistance can be co-dependent, and that in certain cases, defense priming can drive developmental processes and promote plant yield.

**Summary statement:** Growth-defense tradeoffs in plants result in loss of yield. Here, we demonstrate that immunity priming in different pathways uncouples this tradeoff and allows for disease resistant plants with robust growth.

## Introduction

Plants often face abiotic and biotic stresses such as pathogen infection or climate/ ecological changes. These stresses induce physiological, biochemical, and molecular changes, which can be reflected in altered development, lowered growth, and reduced productivity. Plants lack mobile immune cells and an adaptive immune system, and protect themselves against pathogens using several complex mechanisms. In the interaction between the plant and the attacker, elicitor molecules of various types are released (Mishra et al., 2012). Perception of elicitors by the host plant results in signaling that leads to activation of various defense mechanisms (Walters and Heil, 2007).

Following pathogen infection, plants can develop enhanced resistance to further attack. This is known as induced resistance, and is broadly divided into systemic acquired resistance (SAR) and induced systemic resistance (ISR) (Walters and Heil, 2007). SAR, commonly triggered by local infection, can provide long-term resistance to subsequent infection (Klessig et al., 2018). During SAR, pathogenesis-related (PR) genes are activateds. Among the best-characterized PR genes is *PR-1*, which is often used as a marker for SAR (Pieterse et al., 2014). SAR generally relies on salicylic acid (SA), which has been shown to increase in plant tissues during pathogenesis (Yalpani et al., 1991) . The SA receptor NPR1 (Non-expressor of Pathogenesis Related genes 1) regulates SAR signaling following SA perception. NPR1 is activated by SA, and subsequently activates transcription of genes required for SAR, including PR genes (Cao et al., 1997; Chen et al., 2021; Pieterse et al., 2014).

Unlike SAR, which is usually activated during pathogen infection, induced systemic resistance- ISR is usually described as being initiated by beneficial soilborne microorganisms, including rhizobacteria and fungi. ISR is associated with a physiological state in which plants can react more efficiently to pathogen attack (Shoresh et al., 2005), known as immunity “priming”. ISR is regulated by jasmonic acid (JA) and ethylene (ET), and is typically not associated with a direct activation of PR genes, although *NPR1* can enhance ISR and, in some cases, *PR-1* is accumulated during ISR priming (Gupta and Bar, 2020; Meller Harel et al., 2014; Pieterse et al., 1998). As a result of priming, upon secondary infection, defense responses occur more rapidly and strongly than during the primary infection, enabling a more effective response to the new infection (Heil and Bostock, 2002).

While the deployment of defense mechanisms is imperative for plant survival, defense activation can come at the expense of plant growth. This is known as the “growth-defense tradeoff” phenomenon, and is the result of a balance between growth and defense/ adaptation responses (Figueroa-Macías et al., 2021). Growth– defense tradeoffs have an important role in plant survival (Huot et al., 2014). In general, mature organs and tissues are no longer growing and, as a result, can be more prepared for defense as they possess more energy and resources that can be mobilized. One recent work suggested that competition for resources could shape the costs of defense induction (Karasov et al., 2017). Growth-defense trade-offs in plants have been increasingly investigated of late (Ke et al., 2020; Margalha et al., 2019; Monson et al., 2022; Saini and Nandi, 2022). Although some works have suggested that growth and defense are antagonistic, the extent of the “development-defense-tradeoff” and the mechanisms by which it affects plant growth and agricultural yield are largely uninvestigated.

Phytohormones involved in both developmental processes and defense, could potentially be involved in regulating development-defense tradeoffs. The developmental phytohormone Cytokinin (CK) is involved in cell division, leaf senescence, apical dominance, vascular differentiation, root development and stress responses (Zürcher and Müller, 2016). CKs have reported roles in response to biotic stresses (Großkinsky et al., 2011; Gupta et al., 2020; Jiang et al., 2013), and promote resistance through the SA signaling pathway (Choi et al., 2010; Gupta et al., 2020). We have previously shown that CK-deficiency results in higher disease susceptibility, while high endogenous CK content, as well as external application of CK, confer disease resistance (Gupta et al., 2020). Gibberellins (GA) are plant growth hormones involved in stem elongation, germination, leaf expansion, flowering, and fruit development (Davière and Achard, 2013). GAs regulate growth by destabilizing DELLA, a negative regulator of GA signaling (Locascio et al., 2013). GAs can negatively regulate JA signaling, potentially affecting disease resistance (Hou et al., 2010). The central roles of CK and GA in plant growth and development, together with their reported connections with biotic stresses, suggest that these two hormones could be involved in growth- or development-defense tradeoffs.

SA and JA, which were classically defined as “defense/ stress” hormones, also regulate developmental processes (van Butselaar and Van den Ackerveken, 2020; Ghorbel et al., 2021; Rivas-San Vicente and Plasencia, 2011; Wasternack et al., 2013). This suggests that growth and defense may not be simply antagonistic, but rather, have a more complex balance (Koo et al., 2020). Rather than looking at growth and defense as a binary switch, a more holistic view suggests that plant fitness is optimized when growth and defense are appropriately prioritized in response to both environmental and developmental cues. Researching the molecular mechanisms used by plants to balance growth and defense can lead to the possibility of manipulating these mechanisms, with the goal of increasing plant productivity.

In this work, we aimed to investigate the growth-defense trade-off in tomato, by examining how defense priming affects tomato development and agricultural qualities. We used three different immunity elicitors, ASM (Acibenzolar-S-Methyl), also known as BTH (benzo (1,2,3) thiadiazole-7-carboxylic acid) and marketed under the trade name “bion” (Oostendorp et al., 2001), a SAR elicitor, *Trichoderma harzianum*, a classical fungal ISR elicitor (Elad, 2000a), and *Bacillus megaterium*, a bacterial elicitor which likely induces both SAR and ISR (Gupta et al., 2021a, 2022a). These elicitors were employed to investigate differences and commonalities between SAR and ISR pathways, and examine their effect on tomato development. We found that both SAR and ISR affect plant development, with ISR generating a more pronounced positive effect on agricultural qualities. We show that growth and defense can be positively correlated, as mutants that lost the ability to activate defenses, also lost the growth effects. In addition, we found that cytokinin response is activated in response to ISR, and is required for the observed growth and disease resistance resulting from ISR activation. Taken together, our results suggest that growth promotion and induced resistance can be co-dependent, and that defense priming can, within a certain developmental window, not necessitate a trade-off on growth, but rather, encourage and drive developmental processes.

## Results

### Activation of SAR and ISR both promote resistance to B. cinerea and increase plant immunity

The elicitors *Trichoderma harzianum* T39, a *Trichoderma* sp. fungus that induces immunity via the JA/ET pathway (Meller Harel et al., 2014), ASM (Acibenzolar-S- Methyl), a SA analog that primes the SA pathway (Huot et al., 2014), and *B. megaterium* “4C”, a phyllosphere *Bacillus* isolate that promotes plant development and immunity and likely activates both SAR and ISR pathways (Gupta et al., 2021a, 2022a), were used to compare SAR and ISR induction of pathogen resistance.

Elicitors were applied via foliar spray. Further details on immunity priming methodology are provided in the methods section. We conducted *Botrytis cinerea* infection assays on 4-5-week-old primed *S. lycopersicum* M82 plants. We found that pre-treatment with all three elicitors reduced disease levels (**Fig. 1**). Foliar spray or soil-drench application of all three elicitors had a similar effect on disease resistance (**Fig. S1**).

**Figure 1:**
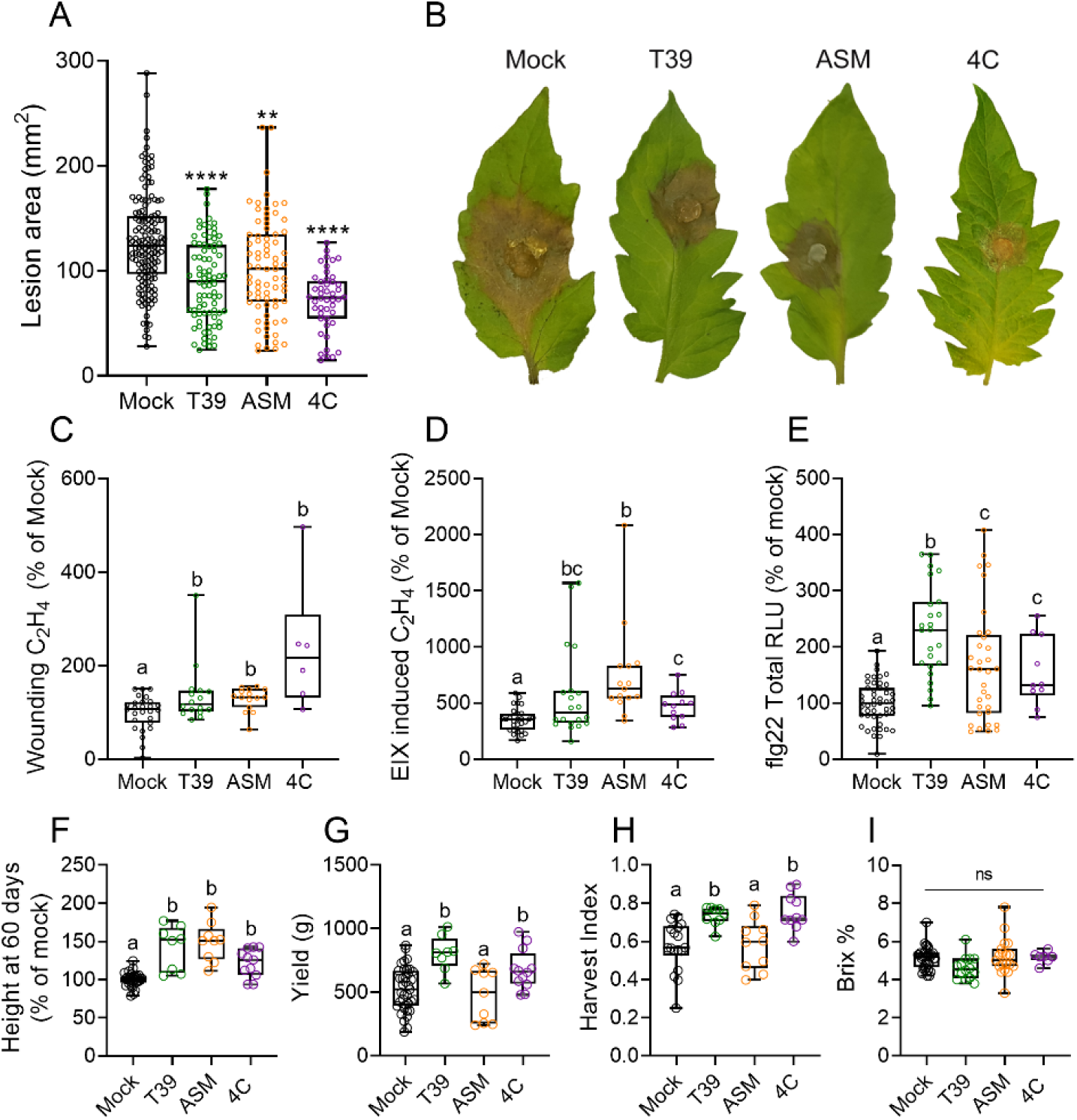
Treatment with T39, ASM, and 4C reduces *B. cinerea* disease, induces immune responses, and promotes plant growth and agricultural parameters. **A-B** *S. lycopersicum* cv M82 4-5-week-old Tomato plants were sprayed twice with elicitors: 3 days and 4 h before inoculation with 3-day-old *B. cinerea* mycelia. Mock- DDW tretment. Lesion area was measured 5 days after inoculation. Experiment was repeated 5 independent times. **A** Lesion area. **B** representative images, 5 days after inoculation. **C-D** Ethylene production was measured alone (C) or with the addition of 1 μg / mL EIX (D), using gas-chromatography. **E** ROS production following flg22 treatment was measured using the HRP luminol method, and expressed in terms of total RLU (Relative Luminescence Units). **F-I** *S. lycopersicum* cv M82 Tomato plants were soil-drenched with elicitors or DDW (Mock) once a week for four weeks. **F** Height of 60-day-old plants. **G** Average fruit weight per plant. **H** Yield per vegetative plant weight, expressed as Harvest Index. **I** Average tomato soluble sugars, expressed as Brix %. Boxplots indicate inner-quartile ranges (box), outer-quartile ranges (whiskers), median (line), all points shown. Asterisks indicate significance from mock treatment (A), and different letters indicate statistically significant differences among samples (C-H) in Welch’s ANOVA with Dunnett’s post hoc test (A, E, F), one-way ANOVA with Tukey’s post hoc test (D, G), or an unpaired two-tailed t-test with Welch’s correction (C, H-I). A: N = 82, \*\**p*<0.01; \*\*\*\**p*<0.0001. C: N=15, p<0.05. D: N>10, p<0.047. E: N>12, p<0.0007. F: N=8, p<0.018. G: N=9, p<0.04. H: N=10, p<0.004. I: N>14, ns=not significant.

Induction of plant immunity was also examined using both SAR and ISR elicitors. We measured ethylene and ROS (reactive oxygen species) production in 4-5-week-old plants. These assays are indices of immunity activation (Leibman-Markus et al., 2017). We found that pre-treatment via foliar spray with the ISR elicitor T39, the SAR elicitor ASM, or the combined SAR/ISR elicitor 4C, all amplified wounding ethylene production and ethylene production mediated by the fungal MAMP (microbe associated molecular pattern) EIX (ethylene inducing xylanase, see (Anand et al., 2021)) in *S. lycopersicum* M82, as compared to mock-treated plants (**Fig. 1C-D**). Pre-treatment with T39, ASM, or 4C, also amplified ROS production in response to the bacterial PAMP (pathogen associated molecular pattern) flg22 (**Fig. 1E**).

### Activation of SAR or ISR differentially promote plant development and productivity

To examine possible growth-defense tradeoffs, we treated plants separately with elicitors, inducing immunity via ISR with T39, via SAR with ASM, or via both pathways, with 4C. ASM concentrations were calibrated in preliminary assays (**Fig. S2**; see methods section for further details). Plants were soil-drenched with the different elicitors once a week for 4 weeks, and grown to harvest, and agricultural parameters were sampled. All three elicitors promoted plant growth (**Fig. 1F**), however, only T39 and 4C were able to significantly promote yield and Harvest index (**Fig. 1G-H**). ASM significantly reduced yield. Total tomato soluble sugars, as measured by Brix refractometry, were unaffected (**Fig. 1I**), suggesting that immunity elicitation did not affect metabolic processes governing fruit quality.

We next examined whether immunity elicitation could affect seedling development (**Fig. 2**). 10-day-old seedlings were spray-treated with elicitors once and harvested after a week for growth and development quantification, or given two treatments one week apart, and harvested one week after the second treatment. All three elicitors affected seedling height (**Fig. 2A, B**), though ASM was only able to do so after two treatments (**Fig. 2B**), while T39 and 4C increased plant height after one treatment (**Fig. 2A**). T39 and 4C also increased seedling fresh weight (**Fig. 2C, D**) and the number of leaves produced by the seedling (**Fig. 2E, F**). ASM did not affect fresh weight or leaf numbers. Representative images of treated seedlings (**Fig. 2 G-J**), meristem and four youngest leaf primordia (**Fig. 2 K-N**), and fifth leaves (**Fig. 2 O-R**) are provided. A more in depth analysis of the effect of T39 on development demonstrated that T39 treatment resulted in increases in the height of the shoot apical meristem (SAM) but not the cotyledons (**Fig. S3A,E,F**), indicating that the treatment affected hypocotyl growth mostly above the SAM. T39 treatment also promoted the number of unfurled leaves (**S3B,G**), the developmental plastochron (**Fig. S3C, H**) and the meristematic differentiation to flowering (**Fig. S3D, I**). We observed that T39 promotes leaf development, increasing the number of leaves produced (**Fig. S4A**), and leaf complexity (**Fig. S4B-D**) (**Fig. S4A, B**). T39 did not alter the developmental program of leaves 1-2, which are simple and have a short developmental window in tomato (Shleizer-Burko et al., 2011) (**Fig. S4C**). We previously reported that 4C treatment promotes the leaf developmental program, increasing leaf complexity and patterning (Gupta et al., 2022a). ASM had no effect on leaf development (**Figs. 2, S4**).

**Figure 2:**
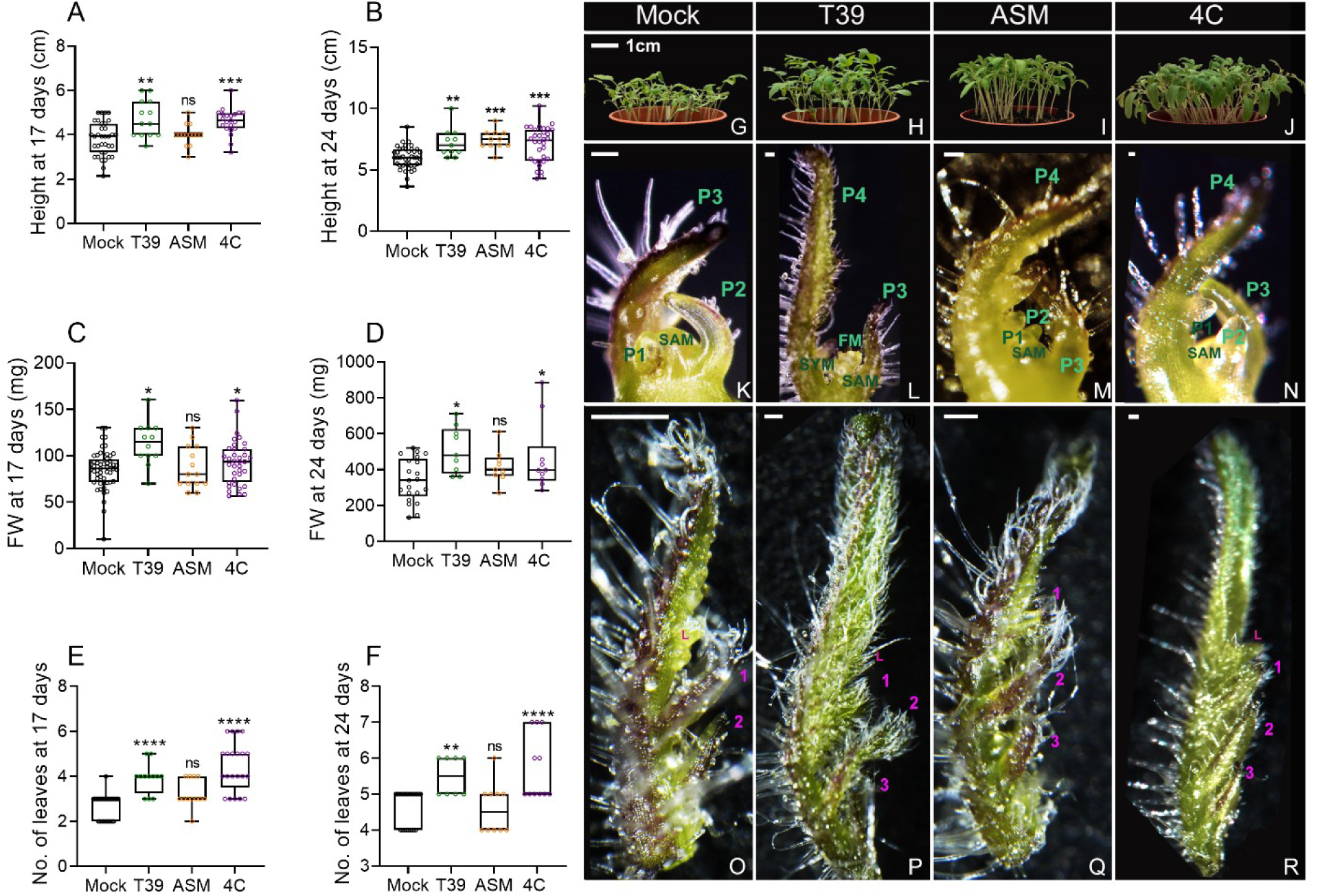
Treatment with T39, ASM, and 4C affects seedling growth and development. *S. lycopersicum* cv M82 Tomato seedlings were sprayed with elicitors or DDW (Mock) once (at 10 days of age, **A,C,E**) or twice (at 10 and 17 days of age, **B,D,F**). **A,B** Height. **C, D** Fresh weight. **E, F** Number of leaves. Boxplots indicate inner-quartile ranges (box), outer-quartile ranges (whiskers), median (line), all points shown. Asterisks indicate statistical significance from Mock treatment in one- way ANOVAwith Bonferroni’sposthoctest, A: N=14, p<0.0003; B: N=12, p<0.0063; C: N=12, p<0.013; D: N=10, p<0.04; E:N=12, p<0.0001, F: N=8, p<0.0047. **G-R** representative images at 24 days of seedlings (**G-J**), a shoot with the two oldest leaves removed (**K-N**), and Leaf No. 5 (**O-R**), which was at the P5 stage in Mock and ASM (**O, Q**), and at the P6 stage in T39 and 4C (**P,R**). **K-R** were acquired using a Nikon SMZ25 stereomicroscope equipped with a Nikon DS-RI2 camera and NIS elements software. Bars: G (For G-J): 1cm, K-N: 100 µM, O-R 250 µM. SAM= shoot apical meristem; FM= floral meristem; SYM= Sympodial meristem (the vegetative meristem that continues to produce leaves after transition to flowering). P1/2/3/4 indicates developmental plastochron. In O-R, numbers indicate pairs of initiated leaflets. “L” indicates lobe.

### The balance between cytokinin and gibberellin is involved in altered development in response to immunity elicitation

We next examined possible routes through which T39, ASM, and 4C promote development. The balance between CK and GA determines the leaf developmental window in tomato (Israeli et al., 2021). As we observed that the leaf developmental window was extended upon T39 and 4C treatment, with additional organs being produced (**Figs. 2, S3-S4,** see also (Gupta et al., 2022a)), we hypothesized that changes in CK and/or GA content or response might be involved. Therefore, we assayed CK and GA pathway gene expression and content, in plant leaves spray- treated with the different elicitors, in both source (**Fig. 3**) and sink (**Fig. S5**) leaves. Additional details are provided in the methods section. In source leaves, CK biosynthesis (*IPT-* isopentenyl transferase genes (Žižková et al., 2015)), activation (*LOG-* lonely guy genes (Naseem et al., 2015)), and degradation (*CKX*- cytokinin dehydrogenase genes (Cueno et al., 2012)) were up-regulated by all elicitors (**Fig. 3A-C**), with some differences. CK degradation appeared more strongly activated by ASM and 4C as compared with T39 (**Fig. 3C**). T39 and 4C treatment resulted in an increase of isopentenyl (iP) in source leaf tissue (**Fig. 3G**), and 4C treatment increased isopentenyl riboside (iPR) content. Despite gene expression results, ASM treatment did not increase CK content, and resulted in reduction in the content of transzeatin (tZ, **Fig. 3G**). Indeed, CK signaling (*TRR*- Tomato Response regulator genes (Bar et al., 2016)) was promoted by T39 and 4C, but not ASM (**Fig. 3D**). *TRRs* are numbered based on their homology to *Arabidopsis ARR*s, such that, for example, *TRR3/4* is homologous to both *ARR3* and *ARR4* of *Arabidopsis*, and so on. GA biosynthesis genes (*GA20 oxidase* and *GA3 oxidase* (Shohat et al., 2021)) were both up and down regulated upon ASM or 4C treatment, while T39 caused a small up-regulation in *GA3ox2* (**Fig. 3E**). The GA degrading *GA2ox7* (Shohat et al., 2021) was up regulated following T39 and ASM treatment, but not 4C treatment (**Fig. 3F**). All three elicitors caused an increase in GA4 and a decrease in GA7 (**Fig. 3G**). Changes in the expression levels of both *GA20ox* and *GA3ox* can explain the shift between these two GA compounds (Hedden, 2020). To confirm defense pathway activation, we assayed the expression of the classical SAR gene *PR1a* (Silverman et al., 2005) and the classical ISR gene *LoxD* (Mariutto et al., 2011), and the ratio between the expression of these two genes was calculated (**Fig. 3H**). While we cannot rule out activation of both pathways, a ratio below one is indicative of ISR being the primary defense pathway, while a ratio above one suggests that SAR is activated. Our results suggest SAR as the primary pathway for ASM, as previously reported (Huot et al., 2014), and ISR as the primary pathway for T39, as previously reported (Meller Harel et al., 2014). The interim, above one value for 4C supports combined ISR and SAR activation with this bacterium, as we previously suggested (Gupta et al., 2021a, 2022a).

**Figure 3:**
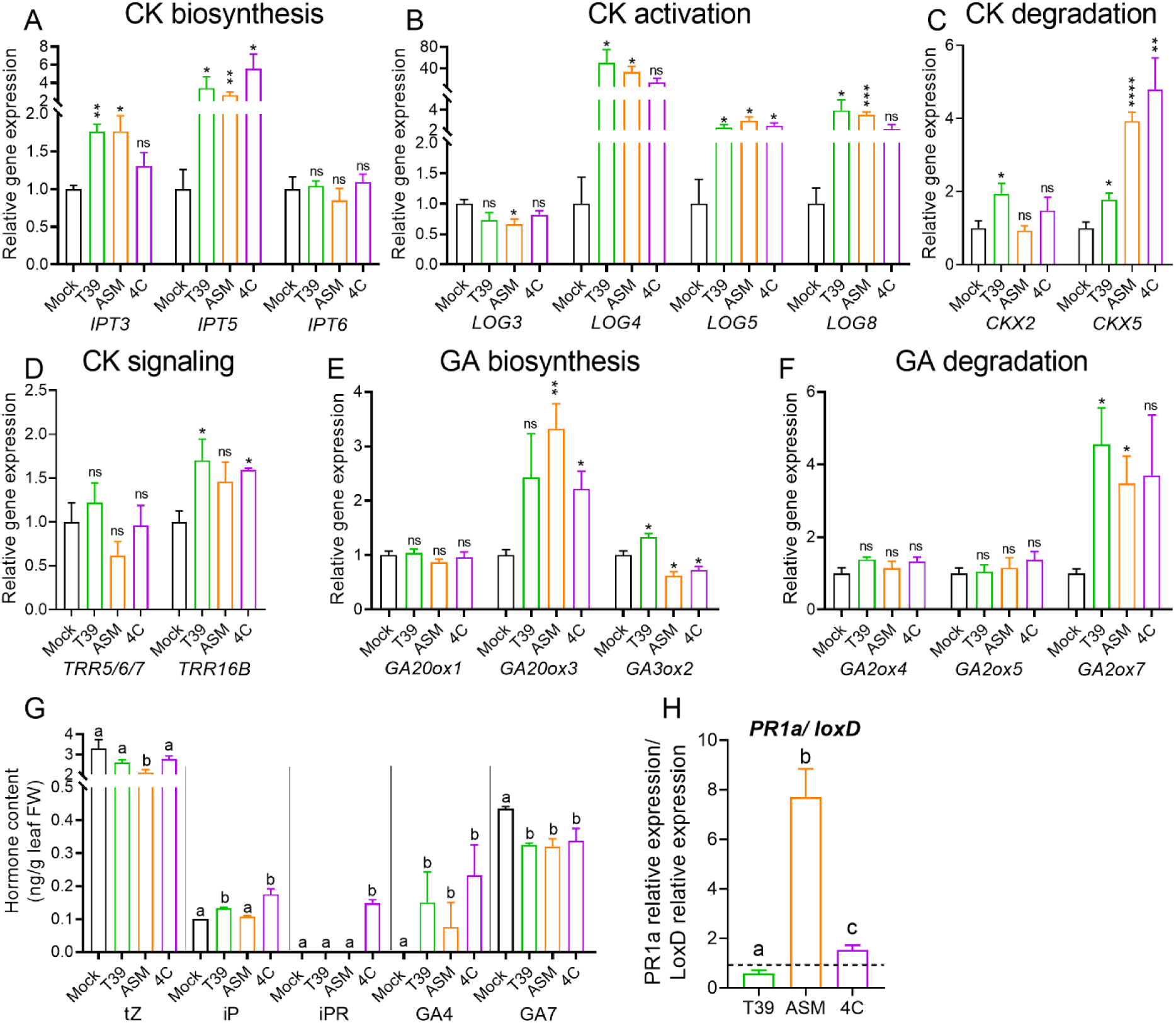
Treatment with elicitors affects CK and GA pathway gene expression and hormone content in source leaves. *S. lycopersicum* cv M82 Tomato seedlings were sprayed with elicitors or DDW (Mock) twice, at 10 and 17 days of age. RNA was prepared from the second and third leaves, 48 h after the second treatment. qRT-PCR relative expression was normalized to the geometric mean of the expression of 3 normalizer genes: *EXP (* Solyc07g025390), *CYP* (Solyc01g111170), and *RPL8* (Solyc10g006580). **A** Cytokinin biosynthesis *IPT*genes. **B** Cytokinin activation *LOG* genes. **C** Cytokinin degradation *CKX* genes. D Cytokinin response regulators (*TRRs*). **E**Gibberellinbiosynthesisgenes(*GA20ox*/ *GA3ox*). **F** Gibberellin degradation genes (*GA2ox*). **G** Quantification of CK and GA derivatives. tZ- transZeatin, iP- isopentenyl, iPR- isopentenylriboside. GA4/7- Gibberellin 4/7. **H** Ratio of the expression of the defense genes *PR1a* and *LoxD*. Below 1 suggests that ISR is the primary pathway activated, above 1 suggests SAR activation. Bars represent mean ±SE. A-F, H: N=5. A-F: Asterisks represent statistical significance from Mock treatment in two-tailed *t*-test with Welch’s correction where appropriate (unequal variances), for each gene. *p<0.05, **p<0.01, ***p<0.001, ****p<0.0001, ns- not significant. G: N=4. G-H: Different letters represent statistically significant differences among samples in a two-tailed *t*-test with Welch’s correction where appropriate (unequal variances). G: p<0.05, H: p<0.0025.

In sink leaves, CK and GA were much harder to detect, likely due to the smaller amount of available tissue. However, alterations to CK and GA pathway genes were also observed in sink leaves (**Fig. S5**). Treatment with 4C activated one *IPT* and one *LOG* gene (**Fig. S5A-B**). All treatments promoted CK degradation (**Fig. S5C**). T39 increased the expression of three *TRR* genes, 4C of two *TRR* genes, and ASM- of one *TRR* gene (**Fig. S5D**). GA biosynthesis was activated by 4C, and repressed by T39 (**Fig. S5E**), and GA degradation was repressed by both T39 and ASM (**Fig. S5F**). CKs and GAs were mostly undetectable in our sink tissues. Transzeatin was unchanged by elicitor treatments, and 4C treatment effected an increase in GA7 (**Fig. S5I**), in accordance with the parallel increase in *GA20ox1* (**Fig. S5E**). Developmental transcription factors, which were not detected in the source leaves, were up- regulated in sink leaves (**Fig. S5G**). The meristem maintenance KNOX gene *TKN2* (Hake and Ori, 2002) and the organ boundary determination gene *GOBLET- GOB* (Berger et al., 2009), were upregulated upon treatment with T39 and 4C (**Fig. S5G**). KNOXI proteins can negatively affect GA levels by repressing the GA biosynthesis gene GA20ox and/or activating the GA inactivation gene GA2ox. Thus, TKN2 could increase the CK/GA ratio by negatively regulating GA levels, which was shown to mediate the function of KNOXI in modulating compound leaf development (Bolduc and Hake, 2009; Hay et al., 2002; Jasinski et al., 2005; Sakamoto et al., 2001). ASM treatment caused up-regulation of *GOB* but not *TKN2*. *Clausa* (*CLAU*), a differentiation promoting transcription factor (Bar et al., 2016), which is known to reduce CK sensitivity and increase GA sensitivity (Bar et al., 2016; Israeli et al., 2021), was unaffected (**Fig. S5G**). The ratio between the expression of *PR1a* and *LoxD* (**Fig. S5H**) was similar to that observed in source leaves (**Fig. 3H**).

The balance between CK and GA, rather that content of each hormone, was reported to be important in leaf development (Fleishon et al., 2011; Israeli et al., 2021). Since we observed increases in CK content and signaling upon elicitor treatments (**Fig.s 3, S5**), we further examined CK signaling in response to elicitors using tomato plants expressing the synthetic cytokinin response promoter *pTCSv2* (two component sensor) driving VENUS fluorescent protein expression (Steiner et al., 2020). Ten-day-old *pTCSv2*-VENUS expressing seedlings received one elicitor treatment, and were imaged after 48 h. All three elicitors increased the CK-response mean fluorescent signal in the meristem and youngest leaf primordia (m+3), while T39 alone increased the mean fluorescent signal in older leaf primordia (P4-P5) (**Fig. 4A-B**). Corrected total fluorescence (CTF), a measure of the total amount of CK- response signal taking into account the fluorescent area and background readings, was unaffected by ASM, but increased in response to T39 and 4C treatment, in meristems and youngest primordia (m+3), as well as in older primordia (P4-P5) (**Fig. 4C**). Thus, T39 and 4C increased both the local signal intensity within the meristem and three youngest leaf primordia, and the total signal measured in all tissues, while ASM increased the local signal intensity in the meristem and three youngest leaf primordia, but did not affect the total CK-response signal (**Fig. 4A-C**; See also (Gupta et al., 2022a)). This correlates with the effect of these elicitors on TRR gene expression in sink tissues (**Fig. S5**), as expected (Steiner et al., 2020; Zürcher et al., 2013).

**Figure 4:**
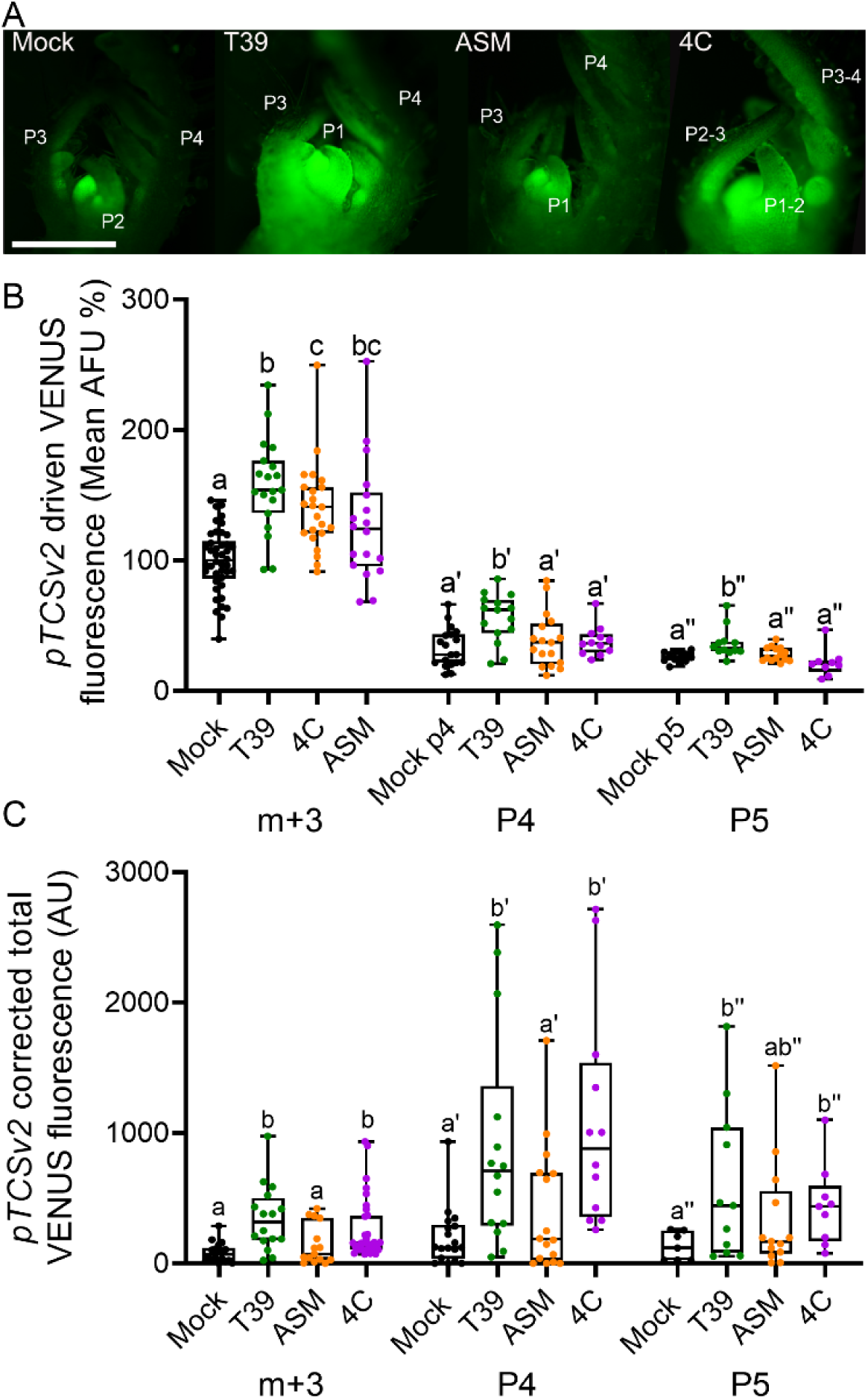
Treatment with T39, ASM, and 4C activates cytokinin response. *S. lycopersicum* cv M82 Tomato seedlings expressing the cytokinin response marker *pTCSv2*::3XVENUSweresprayedwithelicitorsorDDW(Mock)at10daysofage,and imaged after 48 h. **A** Representative shoots. “P” number indicates the developmental plastochron. **B** Mean fluorescent signal, expressed as % of the Mock signal (AFU- arbitrary fluorescent units). m+3= meristem and 3 youngest primordia; P4= fourth plastochron (fourth youngest leaf primordium); P5= fifth plastochron, (fifth youngest leaf primordium). Leaves 1 and 2 were removed, such that the oldest primordium was L3. **C** Corrected total fluorescence (CTF= Integrated Density – (Area of selected cell X Mean fluorescence of background reading), in the indicated tissues. Measurements were taken using FIJI-ImageJ. Boxplots indicate inner-quartile ranges (box), outer-quartile ranges (whiskers), median (line), all points shown. Different letters indicate statistically significant differences among samples in one-way ANOVA with Tukey’s post hoc test or in Welch’s t-test, untagged letters for m+3, tagged letters for P4, double-tagged letters for P5. B: N>9, p<0.048; C: N>12, p<0.05.

Since these results indicated that CK, or the ratio between CK and GA, is required to transduce the developmental signal generated by immunity induction with elicitors, we examined the response to T39, ASM, and 4C treatments, in several altered hormone content and/or signaling lines. Genotypes with high CK content or signaling are generally disease resistant, while genotypes with low CK content or signaling can be disease susceptible (Gupta et al., 2020). Since the ratio between CK and GA can determine developmental outcomes (Israeli et al., 2021), we included both high and low CK, and high and low GA genotypes in the study.

We found that the high CK or low GA lines behaved similar to the M82 background line in growth, development, and *B. cinerea* resistance upon treatment with all three elicitors (**Fig.s 5A-B, C-D, G-H, K-L**, and **S6A-B, C-D, G-H, K-L**). However, T39 and 4C were no longer able to promote growth (**Fig. 5E, I**), development (**Fig. S6E-F, I-J**), or *B. cinerea* resistance (**Fig. 5 F, J)** upon reduced CK (**Fig. 5E, F**) or increased GA signaling (**Fig. 5I, J**). ASM promoted disease resistance irrespective of CK or GA signaling (**Fig. 5B,D,F,H,J,L**), and continued to promote growth upon reduced CK signaling (**Fig. 5E**), but did not affect leaf development in any of the lines (**Figure S6A-L**). Some differences were observed in the level of response of the different lines to the different elicitors (**Fig. S7**), with responses to T39 and 4C significantly lower in *IPT* and *clau* than in M82. Thus, T39 and 4C, which induce ISR, increase CK biosynthesis and response (**Fig.s 3-4**), and cannot promote growth and development in the absence of a functioning CK pathway (**Fig.s 5, S6**). ASM, which induces SAR, does not promote CK biosynthesis (**Fig.s 3**), has minor effects on CK response (**Fig. 4**), does not require a functioning CK pathway to promote growth (**Fig. 5**), and does not promote leaf development (**Fig. S6**). Though 4C induces both SAR and ISR, the ISR effects are dominant in the context of growth and development, and are promoted irrespective of SAR induction.

**Figure 5:**
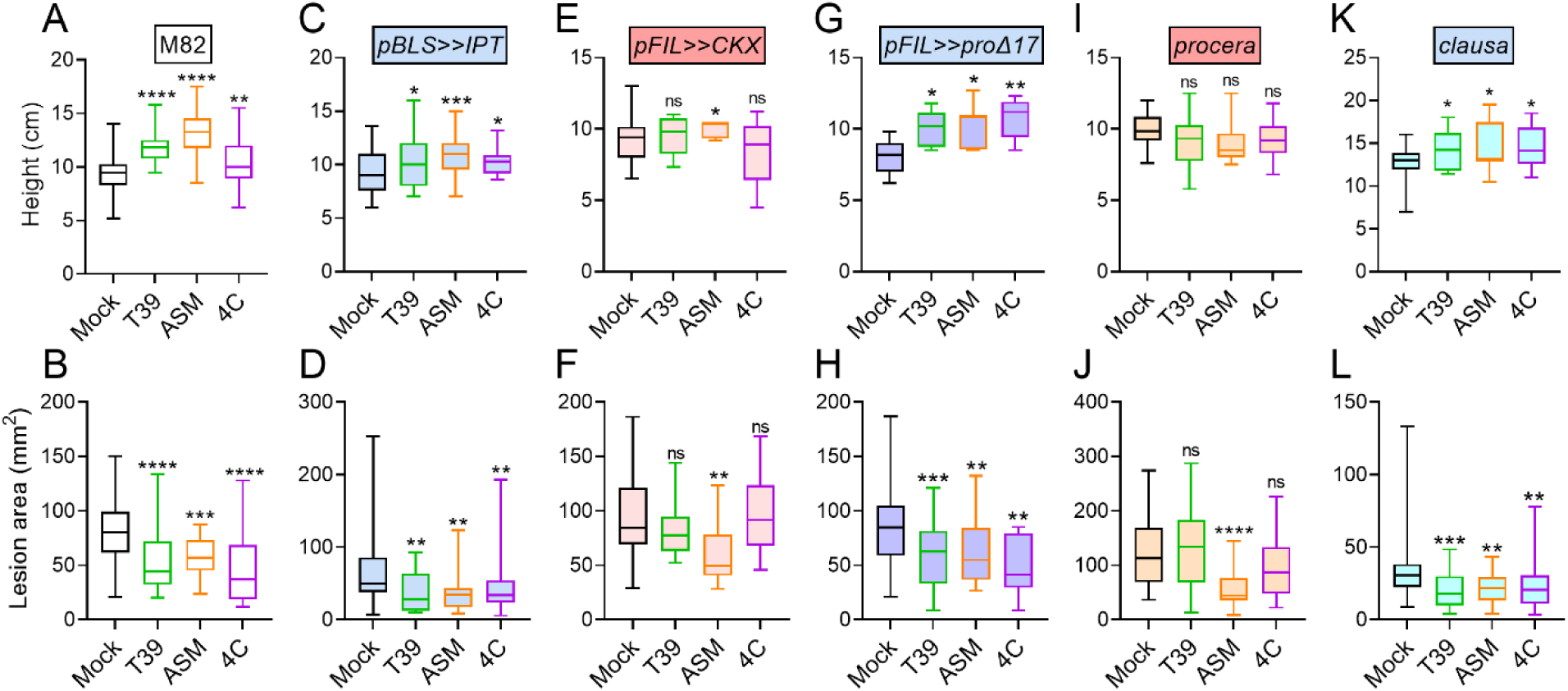
Treatment with T39 and 4C, but not ASM, requires CK to promote growth and disease resistance. **A, C, E, G, I, K** Tomato seedlings of *S. lycopersicum* cv M82 (A-B), the transgenic *pBLS>>IPT* (C-D), *pFIL>>CKX* (E-F), *pFIL>>proΔ17* (G-H), or the recessive mutants *procera* (I-J) or *clausa* (K-L), all in the M82 background, were sprayed with elicitors twice, at 10 and 17 days of age. Seedlings treated with DDW were used as mock. Genotypes with high CK or low GA are indicated in blue, genotypes with low CK or high GA are indicated in red. High/ low refers to content or signaling. See also Table 1 . Height was measured 1 week after the second treatment. **B, D, F, H, J, L** Four-week-old tomato plants of the indicated genotypes were sprayed twice with elicitors or DDW (Mock): 3 days and 4 h before inoculation with 3-day-old *B. cinerea* mycelia. Lesion area was measured 3 days after inoculation. Experiment was repeated 3 independent times. Boxplots indicate inner-quartile ranges (box), outer-quartile ranges (whiskers), median (line), minimum to maximum values. Asterisks indicate statistical significance from Mock treatment in one-way ANOVA with Bonferroni’s post hoc test (A, C, E, G, I, K, J), or in Kruskal-Wallis ANOVA with Dunn’s post hoc test (B, D, F, H, L). A: N=38. B: N=24. C: N=24. D: N=28. E: N=12. F: N=14. G: N=9. H: N=12. I: N=14. J: N=38. K: N=18. L: N=32. *p<0.05, **p<0.01, ***p<0.001, ****p<0.0001, ns- not significant.

**Table 1:**
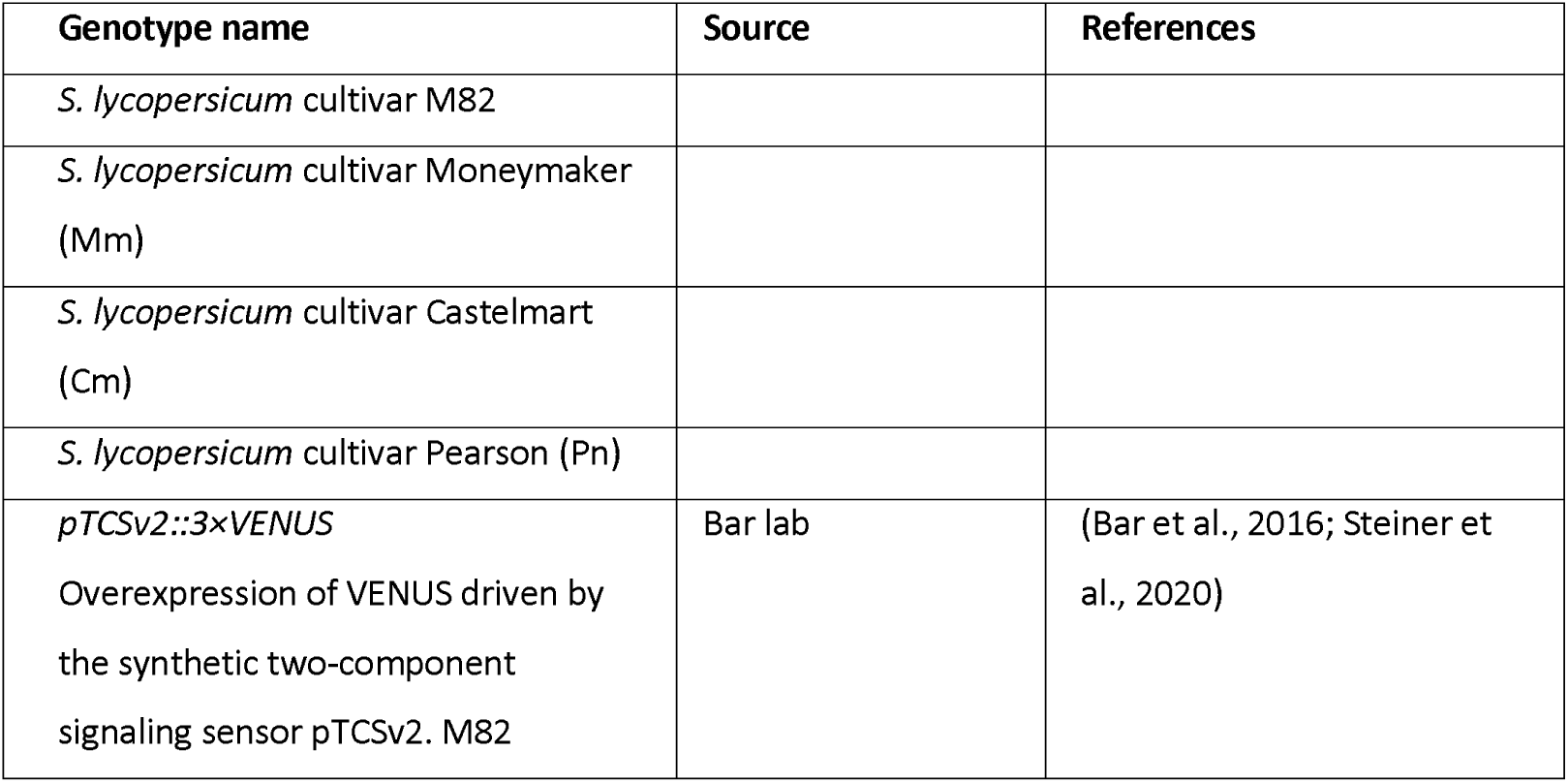
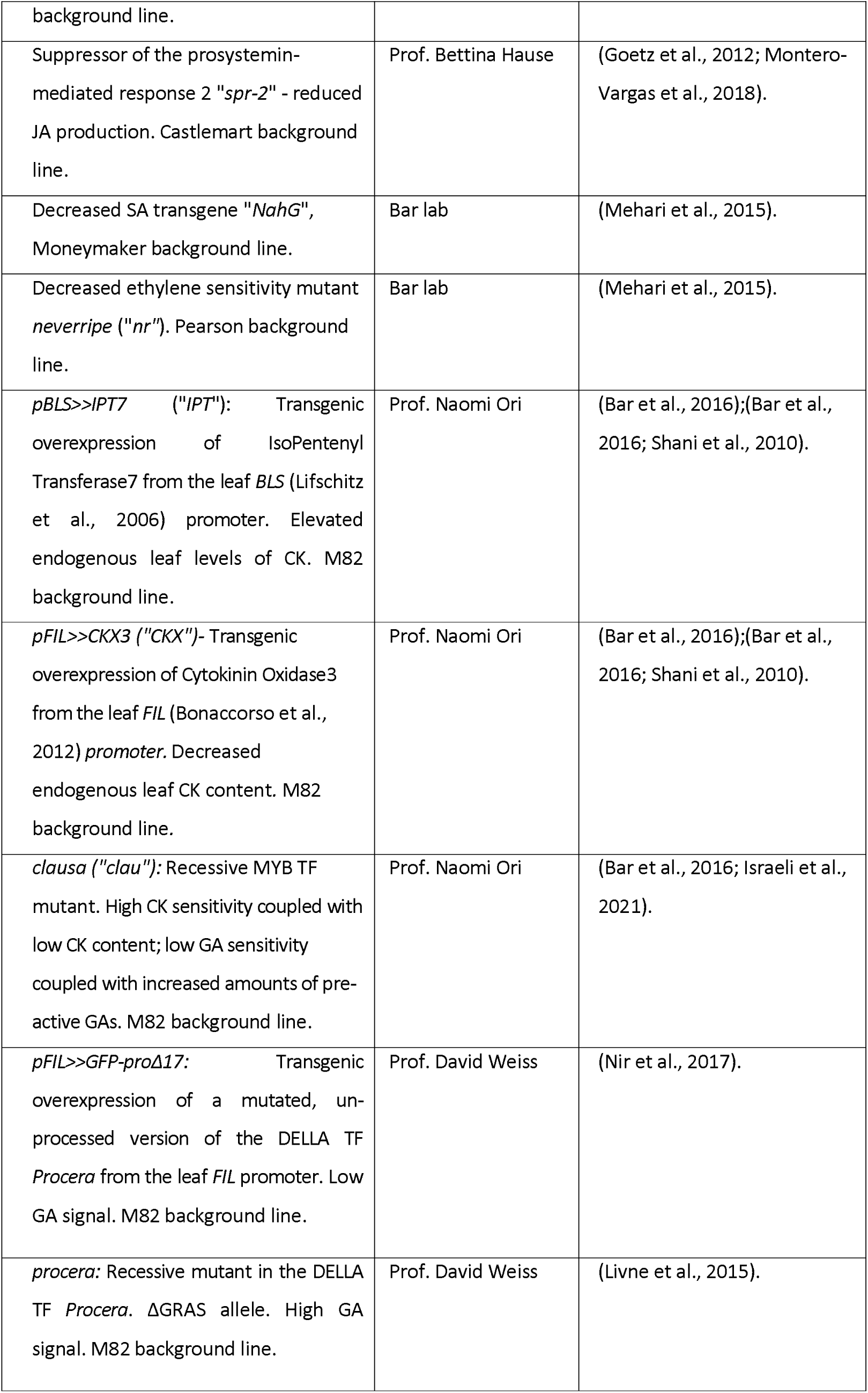
Plant genotypes used in this study.

### Blocking SAR or ISR in tomato mutants leads to loss of disease resistance

To further examine the relationship between development and defense, we used defense mutants. Tomato mutants compromised in jasmonate and ethylene pathways have been previously shown not to respond to *Trichoderma* with disease reduction in certain cases (Jogaiah et al., 2017; Korolev et al., 2008), while lines compromised in the salicylic acid pathway are expected not to respond to ASM (Oostendorp et al., 2001). Thus, we used the SA-lacking transgene *NahG* and its background line Moneymaker (Mm) (Brading et al., 2000), the reduced JA mutant *spr-2* and its background line Castelmart (Cm) (Li et al., 2003), and the reduced ethylene sensitivity mutant *neveripe* (*nr*) and its background line Pearson (Pn) (Lanahan et al., 1994), and examined their disease response to elicitation with T39, ASM, and 4C. The mutant lines responded as expected to *B. cinerea* infection: *spr-2* with increased susceptibility (AbuQamar et al., 2008), and *nr* and *NahG* with similar susceptibility (Mehari et al., 2015; Audenaert et al., 2002). All the background lines responded to the three elicitors with reduction in *B. cinerea* incited disease (**Fig. 6**), as was observed for M82 (**Fig. 1**). However, *NahG* did not respond to ASM or 4C (**Fig. 6A**), while *spr-2* and *nr* did not respond to T39 or 4C (**Fig. 6B-C**).

**Figure 6:**
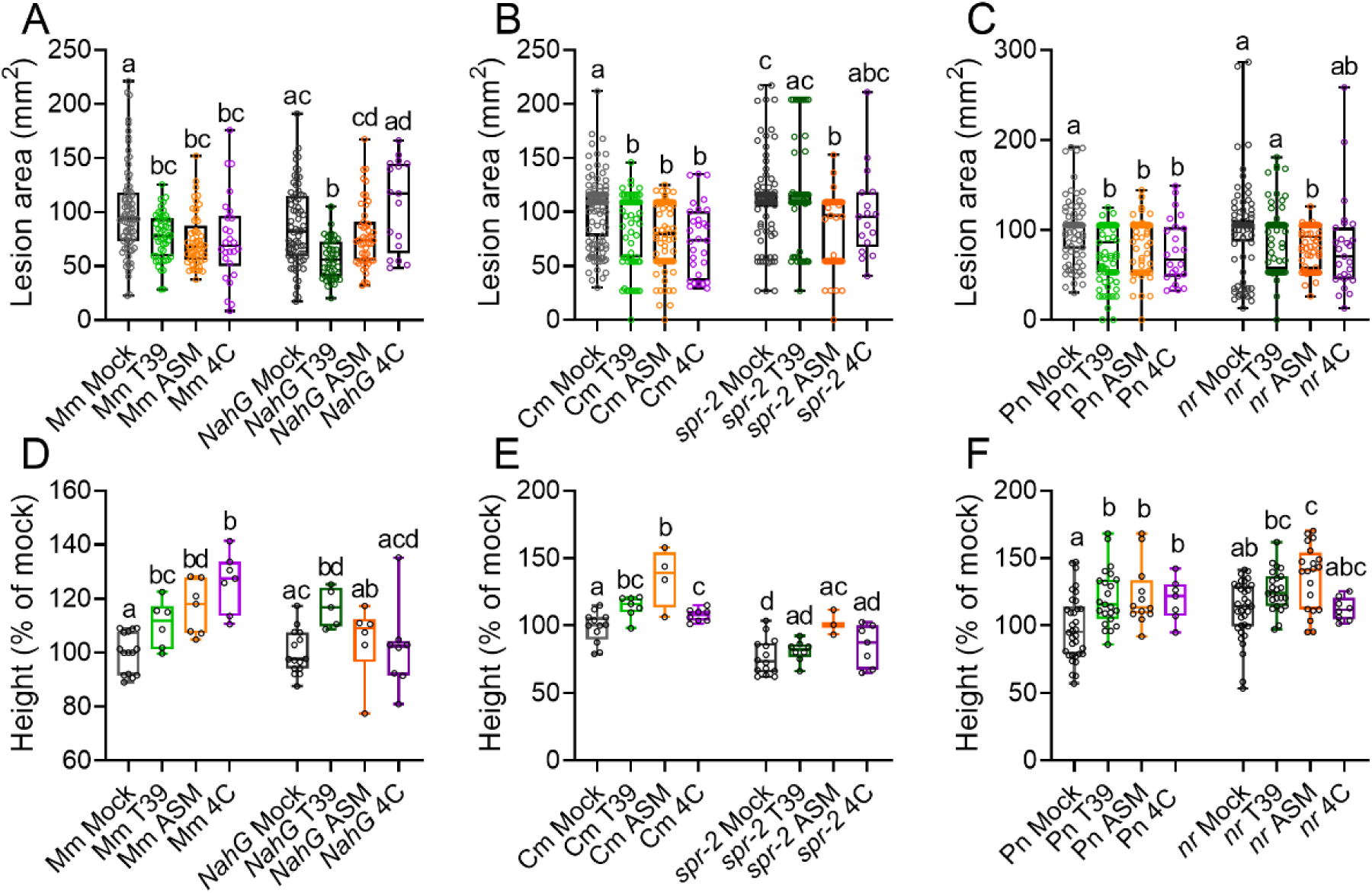
Treatment with T39, ASM, and 4C shows differential effects on disease reduction and development in defense pathway mutants. *S. lycopersicum* plants of the cultivars Moneymaker (Mm) and the reduced SA transgene *NahG* (**A, D**), Castelmart (Cm) and the reduced JA mutant *spr-2* (**B, E**), and Pearson (Pn) and the reduced ET sensitivity mutant *neveripe* (*nr*) (**C, F**), were sprayed with elicitors or DDW (Mock) either twice, at 3 days and 4 h before *Botrytis cinerea* inoculation (A-C), or soil-drenched once a week for four weeks (D-F). **A-C** Plants were inoculated with 3-day-old *B. cinerea* mycelia. Lesion area was measured 3 days after inoculation. Experiment was repeated 4 independent times. **D-F** Height of 60-day-old plants. Boxplots indicate inner-quartile ranges (box), outer-quartile ranges (whiskers), median (line), all points shown. Different letters indicate statistically significant differences among samples in one-way ANOVA with Tukey’s post hoc test (A-C), or Welch’s ANOVA with Dunnett’s post hoc test (D-F). A: N>20, p<0.042. B: N>20, p<0.045. C: N>30, p<0.043. D: N> 5, p<0.045. E: N>3, p<0.042. F: N>12, p<0.035.

### Blocking SAR or ISR in tomato results in loss of developmental promotion and yield increases

Given that mutating defense pathways prevented disease reduction by elicitor pre- treatment, we examined whether the developmental effects of these elicitors would also be abolished in the same mutants. Plants of the different mutant genotypes and their background lines were soil-drenched with elicitors once a week for 4 weeks. All background lines responded to the elicitors with increases in height (**Figure 6D-F**). Once again, *NahG* did not respond to ASM or 4C (**Fig. 6D**), while *spr-2* and *nr* did not respond to T39 or 4C (**Fig. 6E-F**).

We further examined the ability of the elicitor treatments to promote side shoot production and leaf complexity, parameters that are known to be CK-dependent (Mller and Leyser, 2011; Shani et al., 2010; Werner et al., 2001). ASM did not affect either of these parameters in WT lines, but was included as a control in case it might have effects on the mutant lines. T39 and 4C increased side shoot production and leaf complexity in all background lines (**Fig. S8**). While *NahG* responded similarly to its background line Moneymaker in response to T39, it did not respond to 4C (**Fig. S8A, D**). *spr-2* and *nr* did not respond to T39 or 4C (**Fig. S8B-C, E-F**). While *spr-2* had significantly higher side-shoot production than the Cm background line even without elicitor treatment, not all buds were activated, with many plants having only 2-4 activated side-shoots (**Fig. S8B**), therefore, the lack of activity of T39 and 4C on side shoot activation in *spr-2* is not likely to be due to system saturation. ASM had no effect on side shoot or leaflet generation in either WT lines or defense mutants (**Fig. S8**). T39 and 4C promoted yield in all the background lines, but not in *spr-2* or *nr* (**Fig. S9A-C**). T39 was able to promote yield in *NahG,* but 4C was not (**Fig. S9A**). Untreated *NahG* and its background Mm, and *nr* and its background Pn, had similar disease resistance and developmental parameters. *spr-2* baseline disease resistance was lower than its background Cm, and some of the *spr-2* developmental parameters were also altered (**Fig.s 6, S8**).

We also examined seedling development in defense mutant genotypes (**Fig.s 7, S10**). Ten-day-old seedlings were spray-treated with elicitors twice, one week apart, and harvested on day 24, one week after the second treatment. All elicitors promoted seedling height in the background lines (**Fig. 7A-C**). As observed with M82 (**Fig.s 2, S6**), T39 and 4C increased the number of leaves produced (**Fig. S10A-C)**, as well as leaf complexity (**Fig. S10D-F**). As observed with the mature plants (**Fig.s 6, S8-S9**), *NahG* did not respond to ASM or 4C (**Fig. 7A)**, while *spr-2* and *nr* responded to ASM (**Fig. 7B-C**), but did not respond to T39 or 4C (**Fig.s 7B-C, S10B-C, E-F**). In seedlings, untreated *NahG* and its background Mm had similar leaf development, while *nr* and *spr-2* had altered leaf development when compared with their respective background lines Pn and Cm (**Fig. S10**). Untreated seedlings of all mutant lines had similar height as their background lines (**Fig. 7**).

**Figure 7:**
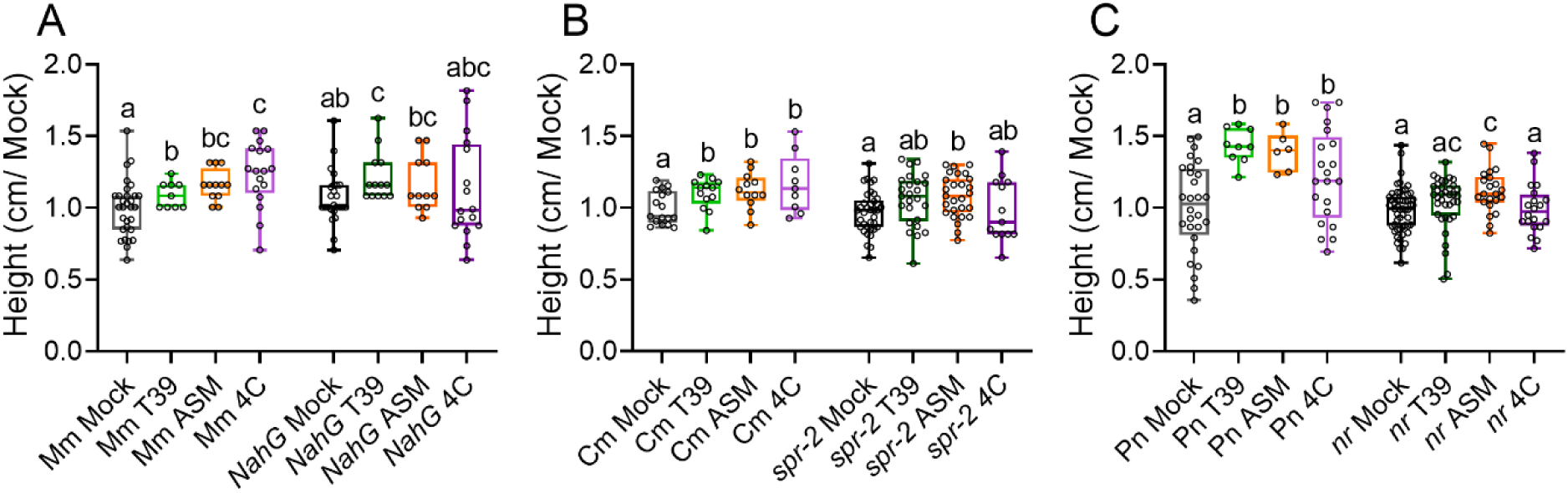
Treatment with T39, ASM, or 4C shows differential effects on plant height in defense pathway mutant seedlings. *S. lycopersicum* seedlingsofthecultivarsMoneymaker(Mm)andthereducedSA transgene NahG (**A**), Castelmart (Cm) and the reduced JA mutant *spr-2* (**B**), and Pearson (Pn) and the reduced ET sensitivity mutant *neveripe* (*nr*) (**C**), were sprayed with elicitors or DDW(Mock)twice,at10and17daysofage.Heightwasmeasured1weekafterthe second treatment. Boxplots indicate inner-quartile ranges (box), outer-quartile ranges (whiskers), median (line), all points shown. Different letters indicate statistically significant differences among samples in one-way ANOVA with Tukey’s post hoc test. A: N>9, p<0.034. B: N>9, p<0.044. C: N>6, p<0.039.

Thus, the competence to respond to each elicitor in increased immunity, is the same as the competence to respond in altered development and increased growth and yield.

### Defense and developmental pathways may have overlapping components

Cellular defense responses are activated by T39 (**Fig. 1**; Gupta et al., 2022b; Gupta and Bar, 2020), ASM (**Fig. 1** ; Marash et al., 2022), and 4C (**Fig. 1** ; Gupta et al., 2021a). ASM treated plants responded to EIX (a JA pathway molecular inducer) more strongly than 4C treated plants, while T39 treated plants responded to flg22 (a SA pathway molecular inducer) more strongly than ASM or 4C treated plants (**Fig. 1**). This could indicate crosstalk or cross-potentiation between SAR and ISR, which was previously suggested (Vlot et al., 2021). To further investigate the relationship between development and defense, we examined whether there could be a genetic overlap between these processes. In previous work (Israeli et al., 2021) we generated a dataset of genes that promote morphogenesis. Our approach, which was validated, relied on published data (Ichihashi et al., 2014) that includes transcriptomics of *S. lycopersicum* cv M82 at four developmental stages. Here, to examine whether we could identify an overlap between morphogenesis and defense pathways, we re-analyzed the previously published data set using KEGG (Kyoto Encyclopedia of Genes and Genomes), to identify additional significantly differential pathways over-represented in morphogenesis. The “plant-pathogen interaction” pathway was found to be significantly enriched in the morphogenetic dataset (Fisher’s exact test, p<0.05). Genes of the plant-pathogen interaction pathway found to be significantly enriched in the morphogenetic group are provided in **Supplemental Table 1** . Defense functions that were found to be upregulated include pattern recognition receptors, mitogen activated protein kinases, WRKY transcription factors, membrane localized NADPH oxidases, calcium-dependent protein kinases, cyclic nucleotide-gated channels, calcium sensors, and nitric oxide synthase. Most of the cellular immune response pathway is represented in the morphogenesis group, suggesting possible genetic overlap between development and defense (see **Supplemental Table1** and **Fig. S11**).

## Discussion

### Activating plant defense does not always generate a growth tradeoff

The concept of growth-defense tradeoff is based on the notion that the limited resources available to an organism must be prioritized according to internal and ambient conditions the organism experiences. In plants, which are sedentary and lack circulatory immunity, the role of ambient conditions, including biotic stresses, becomes more prominent, because plants have limited ability to regulate their environment. Early research of growth defense tradeoffs was partially prompted by observations that plants with highly active defense pathways or defense hormone imbalances can be highly resistant to pathogens, but have severely impaired growth (Huot et al., 2014; Vos et al., 2013). However, many of these reports investigated severely pleiotropic mutants, and dramatic changes to hormonal pathways and hormone levels can have severe impacts on development, particularly considering that defense hormones also have developmental roles (De Vleesschauwer et al., 2014; Vos et al., 2015; Wasternack et al., 2013). Growth-defense tradeoffs have been suggested to rely on crosstalk between different hormones. Cross-talk between CK (Albrecht and Argueso, 2017; Cortleven et al., 2019) or GA (Huot et al., 2014; Monson et al., 2022; Yu et al., 2022), and JA, SA, or ET, has been previously suggested to affect the balance between defense pathways and growth development.

Constitutively active plant defense can result in growth inhibition (Gómez-Gómez et al., 1999; Zipfel et al., 2006), although genetically increasing plant immunity was reported to lack negative effects on plant growth or development in several cases (Cao et al., 1998; Heidel et al., 2004; Leibman-Markus et al., 2021; Pizarro et al., 2020). This suggests that the mode of defense activation, whether by external or genetic means, can affect the level of the “tradeoff”, opening the possibility for activating defense without significantly impairing plant development, a highly sought after agricultural advantage.

### Defense priming can evade negative tradeoffs and result in growth promotion

Previous reports (reviewed in (Gupta and Bar, 2020)) have shown that immunity priming agents can affect plant growth. The underlying hypothesis of the present work was that certain priming agents can in fact improve plant growth, breaking the growth defense tradeoff. To examine this, we selected three known defense priming agents: the SA analog ASM, also known as BTH, a molecular priming agent known to activate SAR (Ishii et al., 2019), the fungus *T. harzianum* T39, a biological priming agent known to activate ISR (Elad, 2000a), and the phyllosphere bacterium *B. megaterium* “4C”, which we previously demonstrated likely activates both SAR and ISR (Gupta et al., 2022a, 2021a).

Our work demonstrates that defense priming can result in both induced resistance and plant growth and development promotion (**Figs. 1-2, 5-7**). T39, ASM, and 4C, all induced resistance against *B. cinerea*. We selected *B. cinerea* as a model pathogen, though this was also reported for all three inducers with additional pathogens (Gupta and Bar 2020; Gupta et al., 2021a; Marolleau et al., 2017;). ASM promoted vertical growth, while T39 and 4C promoted several developmental parameters in both seedlings and mature plants, including agricultural traits (**Fig.s 2,5,S3,S4,S6,S8- S10**). Of note is that treatments with *T. harzianum T39* were previously found to activate resistance to downy mildew in grapevine, without negative effects on plant growth (Palmieri et al., 2012). We previously reported that *B. megaterium* 4C promotes both disease resistance (Gupta et al., 2021a) and plant development and growth, including yield (Gupta et al., 2022a). Though ASM did not positively impact most growth parameters when applied to tomato at 0.001%, it also did not have a negative effect on them in the applied concentration, which could be significant agriculturally under a high disease burden. Higher concentrations of ASM were found to inhibit growth (**Fig. S2**).

### Roles for CK and GA in growth-defense tradeoffs

As central growth hormones, it is perhaps expected that CK and GA should be involved in regulating growth-defense tradeoffs. In recent years, CK has been established as a priming agent, and its role as a defense hormone is increasingly recognized (Choi et al., 2010; Gupta et al., 2020). In the context of disease resistance, most reports indicate that CK promotes immunity and disease resistance through the SA pathway (Choi et al., 2010; Gupta et al., 2020), though roles in JA- mediated defense cannot be ruled out. CK and GA have both previously been suggested to be involved in growth-defense tradeoffs (Cortleven et al., 2019; Yang et al., 2012). CK signaling through response regulators was shown to be regulated following pathogen infection, in several systems (Argueso et al., 2012; Gupta et al., 2020; Igari et al., 2008). CK was suggested to either promote (Choi et al., 2010) – or inhibit (Argueso et al., 2012; Choi et al., 2010) - growth and defense simultaneously, rather than effecting a tradeoff, although experimental evidence of this has thus far been limited.

Notably, JAZ–DELLA interactions integrate JA and GA signaling, such that elevated GA levels or response enhance JAZ repression of defense, while elevated JA levels enhance DELLA-mediated repression of growth (Howe et al., 2018; Yang et al., 2012). Thus, under high GA signaling, plants are more resistant to pathogens that require the SA pathway (Navarro et al., 2008). In many cases, the ratio between CK and GA is the determining factor of the developmental (Israeli et al., 2021) or disease (**Fig. 5**) outcome.

Here, we observed that priming immunity and growth with T39, 4C, or ASM also induced CK signaling (**Figs. 3-4, S5**). CK response was induced more strongly by T39 and 4C than by ASM (**Fig. 4B-C, Fig. 3D, Fig S5D**). CK production was induced by T39 and 4C, but not ASM (**Fig. 3G**). GA biosynthesis and degradation were affected by all elicitors (**Fig. 3**). The promotion of CK signaling by elicitors is unlikely to be explained merely by the changes we observed in hormone levels, and is no doubt a result of increases in CK sensitivity following elicitor application. Notably, the *clausa* mutant possesses significantly reduced levels of many CKs (bar 2016), and increased levels of pre-active GAs (Israeli et al., 2021), but displays increased sensitivity to CK, reduced sensitivity to GA, high-CK developmental phenotypes, and increased disease resistance (Bar et al., 2016; Israeli et al., 2021), demonstrating that hormone sensitivity, and not hormone quantity, determine growth, development, and disease resistance. Thus, the alterations we observed in gene expression following elicitor treatment, can either serve to promote increases in hormone sensitivity, or be the result of feedback relationships aimed at controlling the level of such increases, as previously reported (Müller, 2011; Sun, 2008). Interestingly, ASM promoted plant height and induced disease resistance under CK attenuation, and continued to support disease resistance upon increased GA signaling, while T39 and 4C were not able to promote increases in plant height or disease resistance under CK attenuation or increased GA signaling (**Fig. 5**). Leaf development was promoted by T39 and 4C, but not ASM (**Fig.s 2, S3, S4, S6**). Increased numbers of initiated leaves and increased leaf complexity that resulted from T39 or 4C treatment, could no longer be promoted in low CK or increased GA genotypes (**Fig. S6**). Thus, T39 and 4C treatment can uncouple growth-defense tradeoffs, in a manner dependent on the CK-GA balance. Interestingly, though CK requires SA to induce immunity and promote disease resistance (Choi et al., 2010; Gupta et al., 2020), ASM did not appear to require CK in order to induce immunity and promote vertical growth (**Figure 5E-F**). All three elicitors were found to affect the expression of several assayed CK and GA pathway genes (**Figs. 3, S5**), supporting the notion that activating ISR or SAR results in transcriptional reprogramming that can effect changes in plant developmental programs, and result in increased growth.

### Defense priming competence and plant growth promotion competence have overlapping pathways

SAR depends on SA signaling, while ISR relies on the JA/ET circuit. While SA was shown to be antagonistic to JA in defense signaling (Pieterse et al., 2012, 2009), there are cases where SA and JA were reported to positively interact (De Vleesschauwer et al., 2013; Ferrari et al., 2003; Liu et al., 2016; Ullah et al., 2022). Dependence of ISR on SA signaling has been reported as well (Martínez-Medina et al., 2013). Distinctions between ISR and SAR are not clear cut in tomato, and overlap between the two was reported (Betsuyaku et al., 2018; Liu et al., 2016).

Interestingly, though *Trichoderma* spp. are known to potentiate increases in defense via the JA/ET pathway, T39 activated the SA-dependent *PR1a* (Meller Harel et al., 2014). Furthermore, *B. cinerea* itself, a necrotrophic fungal pathogen against which the plant host requires intact JA signaling to defend itself (AbuQamar et al., 2008), can also activate the SA pathway (Meller Harel et al., 2014). To examine whether the same pathways required for defense in response to priming agents are required for plant growth promotion, we used the SA deficient transgenic line *NahG*, the reduced JA mutant *spr-2*, and the reduced ET sensitivity mutant *neveripe* (*nr*). As expected, ASM was not able to induce resistance in *NahG*, but remained able to do so in *nr* and *spr-2*, while T39 induced resistance in *NahG*, but not in *spr-2* or *nr*. 4C lost the ability to induce disease resistance in all three mutants (**Fig. 6A-C**). Investigating the growth promoting effects in parallel, we observed similar results: T39 promoted growth and increased agricultural and developmental parameters in *NahG*, but was not able to do so in *spr-2* or *nr,* while ASM increased plant height in *spr-2* and *nr*, but not in *NahG*, and 4C could not promote growth, development, or agricultural parameters- in any of the mutant lines (**Fig.s 6D-F, 7, S8-S10**). The background lines behaved as expected. These results support the notion that development-defense tradeoffs can be uncoupled, with priming for defense resistance also serving to promote plant growth, suggesting that components of these two pathways may overlap.

### An overlapping functional “toolbox” is employed in defense and developmental pathways

In a meta-analysis of genes previously defined as morphogenesis promoting (Israeli et al., 2021), the plant-pathogen interaction pathway was found to be up-regulated (**Fig. S11**). While is it tempting to speculate that defense pathways are activated in plant tissues undergoing morphogenesis or growth, another alternative explanation could be that genes with similar functions are required in both defense and developmental processes, resulting in their grouping into similar functional pathways. However, our results indicate that some cross-potentiation between defense and developmental pathways seems likely. We propose that the “development-defense” tradeoff concept should be updated to include the idea that within a defined window, defense priming may actually serve to support developmental processes, rather than inhibit them. **Fig. 8** provides a model conceptualizing the involvement of immunity priming in the mitigation of growth- defense trade-offs. When biotic stress is high, growth is naturally inhibited to free up resources for the activation of stress responses and pathogen resistance mechanisms. Pre-activation of ISR, which primarily requires the JA pathway and promotes CK response, results in rescue of growth inhibition while promoting defense. Pre-activation of SAR, which primarily requires the SA pathway and may not substantially affect the CK/GA balance, promotes defense but results in a more minor growth inhibition rescue as compared with ISR.

**Figure 8:**
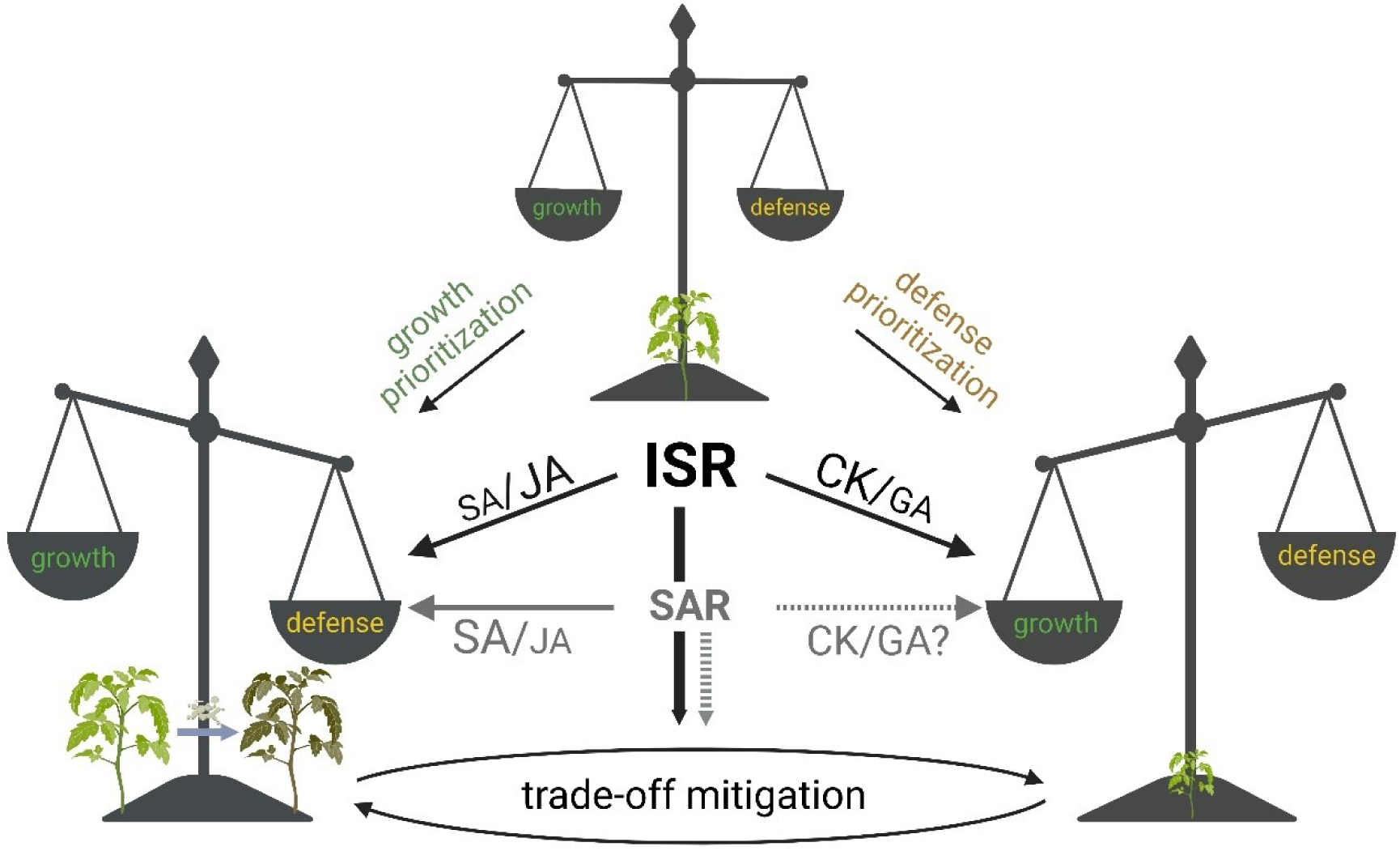
Model conceptualizing the involvement of immunity priming in mitigating growth-defense trade-offs. Plants balance growth/development and defense throughout their lives. When biotic stress is high, growth is naturally inhibited as a mechanism to control resource allocation and better activate immunity and pathogen resistance mechanisms. Pre-activation of ISR, which primarily requires the JA pathway and promotes CK response, results in rescue of growth inhibition while promoting defense. Pre-activation of SAR, which primarily requires the SA pathway and may not substantially affect the CK/GA balance, promotes defense but results in a more minor growth inhibition rescue as compared with ISR. Growth indicated in green, defense indicated in brown/yellow. Illustration generated with Biorender.com. ISR Induced Systemic Resistance SAR Systemic Acquired Resistance SA Salicylic Acid JA Jasmonic Acid CK Cytokinin GA Gibberellin

While plants in nature tend to be uniquely adapted to their particular habitat, and can develop and complete their life cycle in the face of environmental and pathogenic stressors, changing climate and intense cropping practices have in some cases rendered such adaptations ineffective. Our work provides a framework for further research towards the prevention of negative effects that increased immunity can have on plant development and growth. Pending future research, this could potentially be exploited agriculturally in certain cases.

## Materials and methods

### Plant materials and growth conditions

Tomato cultivars and mutant and transgenic lines were used throughout the study as detailed in Table 1. Plants were grown from seeds in soil (Green Mix; Even-Ari, Ashdod, Israel) in a growth chamber (seedling assays) under long day conditions (16 hr:8 hr, light: dark) at 24°C. Mature plants for disease and immunity assays and yield experiments were grown in a 2mm nylon mesh nethouse (spring, summer, and fall- ambient conditions) or in a glasshouse (winter, ambient light supplemented with heating to 24°C).

*pBLS>>IPT*7, “*IPT*”, overexpresses the rate limiting enzyme in biosynthesis of the CK isopentenyl driven from the leaf-specific *BLS* promoter (Lifschitz et al., 2006), and has high CK levels (Márquez-López et al., 2019; Redig et al., 1996; Smigocki and Owens, 1988). *pFIL>>CKX3, “CKX”*, overexpresses a CK degrading dehydrogenase enzyme driven from the leaf specific *FIL* promoter (Bonaccorso et al., 2012), and has low CK levels (Nishiyama et al., 2011; Reid et al., 2016). These lines were previously characterized (Shani et al., 2010). *pFIL>>GFP-proΔ17* “ *proΔ17*” overexpresses a mutated DELLA protein that is constitutively active, driven from the *FIL* promoter, resulting in inhibition of GA signaling (Nir et al., 2017). *procera* “*pro*” is a recessive mutation in the single tomato *DELLA* gene *procera*, resulting in constitutive high GA signaling (Livne et al., 2015). *clausa* “ *clau*” is a recessive MYB transcription factor mutant that has increased CK and reduced GA sensitivity (Israeli et al., 2021).

Promoter driver lines were selected to minimize pleiotropic effects, therefore, *pBLS>>IPT* was used rather than *pFIL>>IPT*. The leaf developmental phonotypes of *CKX* and *pro*, or *IPT*, *clau*, and *proΔ17* are relatively similar in terms of leaf complexity (**Fig. S6**), suggesting that high CK and low GA, or low CK and high GA, can have similar developmental effects, supporting the notion that the ratio between these hormones, rather than their individual content, determine developmental programs.

### Elicitor treatments

For activation of SAR we used ASM (Marolleau et al., 2017), and for activation of ISR we used *T. harzianum* isolate T39 (Elad, 2000b). *B. megaterium* 4C was considered to likely activate both pathways (Gupta et al., 2021a). For disease and immunity assays, elicitors were applied to 4-5-week-old plants by spraying the fourth and fifth leaves to drip. Two treatments were given: 3 days before challenge, and 4 hours before challenge. For agricultural parameters, elicitors were applied to 3-4-week- old plants by soil drench, using 5 mL / pot of the indicated concentrations.

Treatments were applied once a week for four consecutive weeks. For developmental analyses, gene expression and hormone content quantification, elicitors were sprayed to drip on seedlings of the indicated ages. Two treatments were given, one week apart, in all cases except for TCS visualization, in which one treatment was applied.

ASM (Acibenzolar-S-Methyl), also known as BTH (Benzothiadiazole), a synthetic analog of SA, is used commercially to enhance disease resistance (and marketed under the trade name “Bion” (Marolleau et al., 2017; Oostendorp et al., 2001). Previously, application of BTH reduced plant biomass and was correlated with induction of SA-mediated defense responses (Huot et al., 2014). SA analogues have been found to effectively reduce gray mold (*Botrytis cinerea*) disease in tomato (Meller Harel et al., 2014). ASM was shown to be processed in vivo by an SA processing enzyme (Tripathi et al., 2010). As ASM is known to be phytotoxic (Ishii et al., 2019), to work with non-phytotoxic concentrations, we calibrated the system using two different ASM concentrations (**Fig. S2**). 0.01% of ASM were found to reduce seedling height and number of leaves produced (**Fig. S2A-B**). In 4-week-old plants, 0.01% ASM reduced the number of side-shoots produced, but did not negatively affect other plant growth parameters. Treatment with 0.001% of ASM had no negative effects on plant growth (**Fig S2A-B, D-G**), promoted height in both seedlings and 4-week-old plants (**Fig. S2A,D**), and also induced immunity to *B. cinerea*, while 0.01% did not (**Fig. S2C**). Following calibration and verification of lack of toxicity (**Fig. S2**), ASM was applied to plants at a concentration of 0.001% v/v.

*Trichoderma* spp. are established plant growth promoting fungi that enhance disease resistance by inducing ISR in a JA/ET dependent manner (Meller Harel et al., 2014). *Trichoderma* spp. have been shown to promote plant growth and activate ISR against a broad spectrum of pathogens. They are ubiquitous filamentous fungi that colonize the rhizosphere and phyllosphere, promote plant growth, and antagonize numerous foliar and root pathogens (Gupta and Bar, 2020). *T. harzianum* isolate T39 was cultured on PDA plates (Difco lab) at 25°C in natural light. Conidia were harvested in water and applied to plants at a concentration of 10^7^ conidia/ mL.

*Bacilli* spp. are known to promote growth and induce immunity (Miljaković et al., 2020). *B. megaterium* 4C is a phyllosphere isolate we previously demonstrated induces immunity and promotes growth and development of mature plants and seedlings (Gupta et al., 2022a, 2021a). Overnight (Luria-Bertani medium) cultures of *B. megaterium 4C* were washed twice in sterile distilled water, and re-suspended in DW. The cell suspension was adjusted to an optical density of OD _600_=0.1 (approximately equal to 10 ^8^ CFU/ mL), and applied to plants via either foliar spray or soil drench as indicated.

### Measuring growth and agricultural parameters

Plants were treated with T39 or ASM once a week by soil drench for 4 weeks, starting from when the plants reached three unfurled leaves, typically 10-14 days after germination. The first three treatments were given at a volume of 5 ml, and at last treatment was given at a volume of 10 ml. Growth parameters were measured once a week: shoot length was measured using a hand ruler from root crown to shoot apical meristem. The number of side shoots was counted taking into account only shoots of at least 5 cm in length. For leaf complexity, we counted the primary, secondary and intercalary leaflets on the indicated leaf. Primary leaflets are separated by a rachis, and some of them develop secondary and tertiary leaflets. Intercalary leaflets are lateral leaflets that develop from the rachis later than the primary leaflets and between them, and typically have a very short petiolule. The following parameters were measured for yield: weight of fruits, weight of shoot, calculated harvest index (yield weight/total plant weight (shoot + fruit weight), and fruit soluble sugar content, measured as Brix %, using a ref-85 Refractometer (MRC).

### Developmental analysis of seedlings

Seedlings were sprayed to drip with elicitors ten days after germination, and 17 days after germination. Growth and development parameters were analyzed one week after each treatment. Parameters analyzed include height, fresh weight, number of unfurled leaves, leaf developmental stage (No. of the leaf that is Plastochron 1), formation of floral meristem, and leaf complexity. Shoots were dissected with a surgical blade and No.5 forceps. Analyses following dissection were conducted using a Nikon SMZ-25 binocular stereomicroscope or a Dino-Lite hand held microscope.

### *B. cinerea* infection and disease monitoring

Pathogenesis assays were conducted on detached leaves, with the exception of the assays conducted with the defense mutants and their background lines (**Fig. 6**), in which whole plants were inoculated. We have previously demonstrated that in most cases in tomato, relative disease levels are similar on detached leaves and whole plants, though the disease develops faster in detached leaves (Gupta et al., 2020). In the case of the defense mutants, assays were conducted on whole plants, since the similarity of detached leaf and whole plant assays has not been verified in all the employed mutant genotypes. *B. cinerea* isolate *Bc* I16 cultures were maintained on potato dextrose agar (PDA) (Difco Lab) plates and incubated at 22 °C for 5-7 days. 0.4 cm diameter discs were pierced from 5 day old mycelia plates and used for inoculation of both leaves and whole plants. Leaves and plants were kept in a humid chamber following inoculation . The area of the necrotic lesions was measured 3-5 days post inoculation using ImageJ software.

### Ethylene measurement

Ethylene production was measured as previously described (Leibman-Markus et al., 2017). Leaf discs 0.9 cm in diameter were harvested from plants treated as indicated, and average weight was measured for each plant. Discs were washed in water for 1-2 h. Every six discs were sealed with rubber septa in 10-mL Erlenmeyer flasks, containing 1 ml assay buffer (10 mM MES pH 6.0, 250 mM sorbitol) with or without the fungal elicitor ethylene inducing xylanase (EIX), 1 µg/mL, overnight at room temperature. Disposable syringes (5 mL) with 21 gage needles were used to extract the gaseous sample, and rubber stoppers were used to hold the sample until it was injected into the Gas chromatograph. Ethylene content was measured by gas chromatography (Varian 3350, Varian, California, USA).

### Measurement of ROS generation

ROS was measured as previously described (Leibman-Markus et al., 2017). Leaf discs 0.5 cm in diameter were harvested from indicated treatments. Discs were floated in a white 96-well plate (SPL Life Sciences, Korea) containing 250μl distilled water for 24 h at room temperature. After incubation, water was removed, and 50 μL ddH2O water were immediately added to each well to prevent the tissue from drying up. The ROS measurement reaction contained: Luminol 150 μM, HRP 15 μg/mL and either 1lllμM flg22 or 1μg/ mL EIX. Light emission was measured immediately and over the indicated time using a microplate reader luminometer (Spark, Tecan, Switzerland).

### Imaging of the CK-response synthetic promoter *pTCSv2*::3×VENUS

Stable transgenic M82 tomato *pTCSv2*::3 × VENUS plants that express VENUS driven by the synthetic two-component signaling sensor pTCSv2 (Bar et al., 2016; Steiner et al., 2020) were sprayed with elicitors ten days after germination. VENUS expression was analyzed 48 h after treatment using a Nikon SMZ-25 stereomicroscope equipped with a Nikon-D2 camera and NIS Elements v. 5.11 software. ImageJ software was used for analysis and quantification of captured images.

### RNA extraction and qRT-PCR

Plants were sprayed with elicitors twice: ten days after germination, and 1 week later. Plant tissue was collected 48 hours after each treatment. For source leaves, L2 and L3 were harvested from four different plants for each biological replicate. 16 plants were sampled in total. For sink tissue, ∼70 mg of shoot apices (typically consisting of the meristem and 5 youngest leaf primordia, m+5) were collected. 5-7 shoots were sampled per biological replicate, for a total of at least 40 plants. Total RNA was extracted from tissues using Tri-reagent (Sigma-Aldrich) according to the manufacturer’s instructions. RNA (3μg) was converted to first strand cDNA using reverse transcriptase (Promega, United States) and oligo-dT. qRT-PCR was performed according to the Power SYBR Green Master Mix protocol (Life Technologies, Thermo Fisher, United States), using a Rotor-Gene Q machine (Qiagen). Relative expression was quantified by dividing the expression of the relevant gene by the geometric mean of the expression of three established normalizers: ribosomal protein *RPL8* (Solyc10g006580), Cyclophillin *CYP* (Solyc01g111170) (Mascia et al., 2010; Mehari et al., 2015), and EXPRESSED *EXP* (Solyc07g025390) (Bar et al., 2016). These genes were previously demonstrated to possess stable expression levels in immunity or disease challenged tomato plants (Gupta et al., 2021b; Lacerda et al., 2015; Mascia et al., 2010; Mehari et al., 2015; Pizarro et al., 2020). Although the used normalizers are well established, we employed three normalizers to prevent possible effects of lack of stability of any one gene. The geometric mean, rather than arithmetic mean, of the expression levels of these genes was used, in order to account for the possibility that their expression levels are not independent of each other. All primers pairs had efficiencies in the range of 0.97-1.03. Primers used for qRT-PCR are detailed in **Supplemental Table 2**. The genes presented in Figures 3 and S4 are part of a larger subset of genes tested. The full set of tested genes, and the outcome, are provided in **Supplemental Table 3.**

### Quantification of CK and GA in leaf tissues

*S. lycopersicum* cv M82 Tomato seedlings were sprayed with elicitors twice, at 10 and 17 days of age. 48 h after the second treatment, source leaves (L2-L3) and sink leaves (shoot apices) were harvested. Hormone extraction was performed according to (Shaya et al., 2019). Briefly, frozen tissue was ground to a fine powder using a mortar and pestle. One gram of the powder was transferred to a 2 mL tube containing 1 mL extraction solvent (ES) mixture (79% IPA: 20% MeOH: 1% acetic acid) supplemented with 20ng of each deuterium-labelled internal standard (IS, Olomouc, Czech Republic). The tubes were incubated for at 4°C 60min, with rapid shaking, and then centrifuged at 14,000g for 15min at 4°C. The supernatant was collected and transferred to 2mL tubes. 0.5mL of ES was added to the pellet and the extraction steps were repeated twice. The combined extracts were evaporated using speed-vac. Dried samples were dissolved in 200 µl of 50% methanol and, filtered through a 0.22 µm syringe filter with a cellulose membrane. 5‒10 µL were injected for each analysis. LC–MS-MS analyses were conducted using a UPLC-Triple Quadrupole MS (WatersXevo TQMS). Separation was performed on a WatersAcuity UPLC BEH C18 1.7 µm 2.1x100mm column, with a VanGuard pre-column (BEH C18 1.7 um 2.1x5mm). The mobile phase consisted of water (phase A) and acetonitrile (phase B), both containing 0.1% formic acid in the gradient elution mode. The flow rate was 0.3 ml/min, and the column temperature was kept at 35 °C. Acquisition of LC–MS data was performed using MassLynx V4.1 software (Waters). Quantification was done using the isotope-labeled internal standards (IS).

### Data analysis

Data is presented as minimum to maximum values in boxplots, or as average ±SEM in bar graphs. For Gaussian distributed samples, we analyzed the statistical significance of differences between two groups using a two-tailed *t*-test, with additional post hoc correction where appropriate: Welch’s correction for *t*-tests between samples with unequal variances, or Holm-Sidak’s correction when multiple *t*-tests were applied to a data set. We analyzed the statistical significance of differences among three or more groups using analysis of variance (ANOVA).

Regular ANOVA was used for groups with equal variances, and Welch’s ANOVA for groups with unequal variances. Significance in differences between the means of different samples in a group of three or more samples was evaluated using a post- hoc test. Tukey’s post-hoc test was used for samples with equal variances, when the mean of each sample was compared to the mean of every other sample.

Bonferroni’s post-hoc test was used for samples with equal variances, when the mean of each sample was compared to the mean of a control sample. Dunnett’s post-hoc test was used for samples with unequal variances. For samples with non-Gaussian distribution, we analyzed the statistical significance of differences between two groups using a Mann-Whitney U test, and the statistical significance of differences among three or more groups using Kruskal-Wallis ANOVA, with Dunn’s multiple comparison post-hoc test as indicated. Gaussian distribution or lack thereof was determined using the Shapiro-Wilk test for normality. Statistical analyses were conducted using Prism9^TM^.

## Supporting information

All Supplemental materials in one file

## Acknowledgements

We thank Naomi Ori for helpful discussions, and members of the Elad and Bar groups for continuous support.

## Author contributions

M.B. conceived and designed the study. A.S., M.L-M., R.G., I.M., D.R-D., Y.E. and M.B. formulated the methodology and carried out the experiments. A.S., M.L-M., D.R-D., R.G., Y.E. and M.B. analyzed the data. All authors contributed to the writing of the manuscript.

## Data Availability Statement

The authors declare that the data supporting the findings of this study are available within the paper and its Supporting Information files. Raw data is available from the corresponding author upon reasonable request.

