## Supplementary material for "Immunity priming uncouples the growth-defense tradeoff in tomato": All Supplemental materials in one file

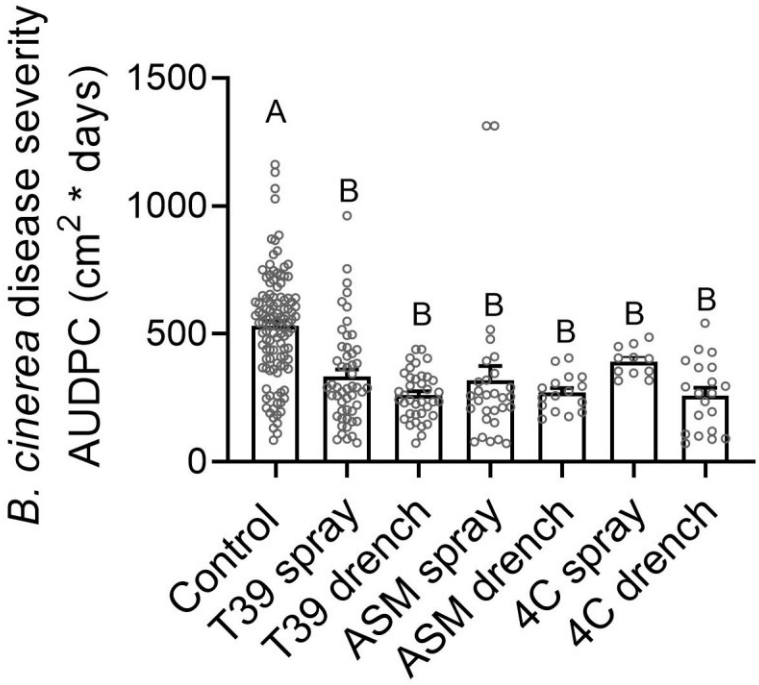


**Figure S1- spray and soil drench application of elicitors results in similar levels of disease reduction.**

Four week old S. lycopersicum M82 plants were sprayed or soil-drenched twice, as indicated, with elicitors: 3 days and 4 h before *Botrytis cinerea* inoculation. Plant were inoculated with 3 day-old *B. cinerea* mycelia. Plants treated with DDW were used as Control. Lesion area was measured 3 days after inoculation. Experiment was repeated 3 independent times.

Bars represent mean ±SE, all points shown. Different letters indicate statistically significant differences between samples in one-way ANOVA with Tukey's post hoc test. N≥12, $p<0.005$.

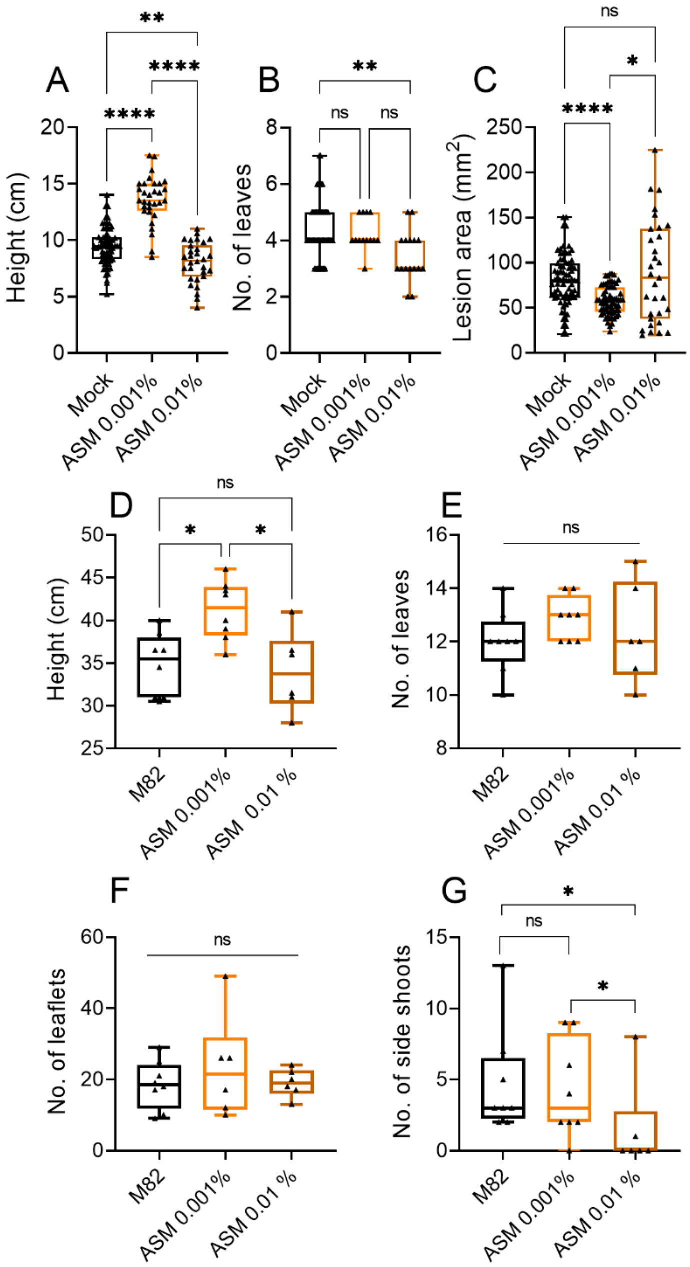


**Figure S2: ASM is not phytotoxic to tomato when applied at 0.001%.**

**A-B** Tomato seedlings of *S. lycopersicum* cv M82 were sprayed with indicated concentrations of ASM twice, at 10 and 17 days of age. Seedlings treated with DDW were used as mock.

**A, B** Height and the number of unfurled leaves were measured 1 week after the second treatment.

**C** Four-week-old plants were sprayed twice with indicated concentrations of ASM: 3 days and 4 h before *Botrytis cinerea* inoculation. Plant were inoculated with 3 day-old *B. cinerea* mycelia. Plants treated with DDW were used as mock. Lesion area was measured 3 days after inoculation. Experiment was repeated three independent times.

**D-F** 3-4-week-old plants were soil-drenched with indicated concentrations of ASM once a week for four weeks. Parameters were measured at 50 days of age.

Boxplots are shown with inter-quartile-ranges (box), medians (line in box) and outer quartile whiskers, minimum to maximum values, all points shown. Asterisks indicate statistical significance among indicated samples in one-way ANOVA with Tukey's post hoc test, A: N=30. B: N=16. C: N=60. D-F: N>6.*$p<0.05$, **$p<0.01$, ****$p<0.0001$, ns- not significant.

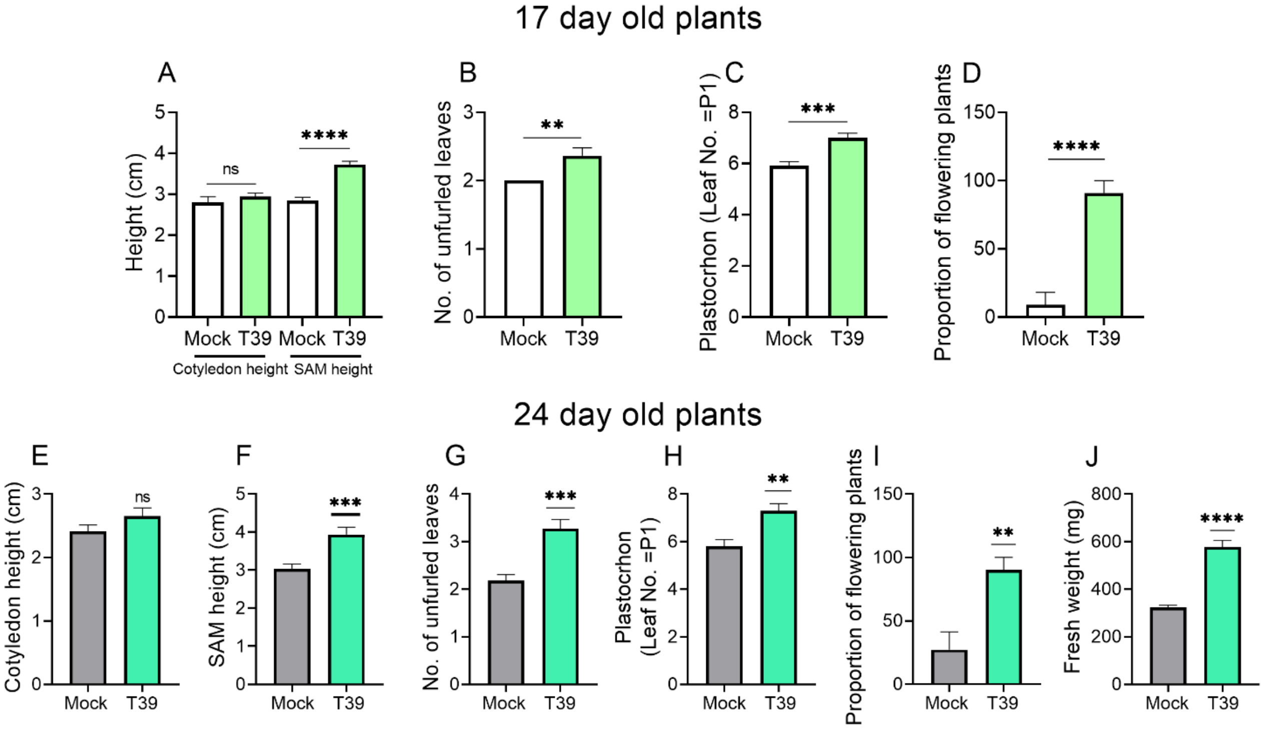


**Figure S3: Treatment with T39 and ASM affects seedling growth and development.**

*S. lycopersicum* cv M82 Tomato seedlings were sprayed with elicitors once (at 10 days of age, **A-D**) or twice (at 10 and 17 days of age, **E-J**). Seedlings treated with DDW were used as mock. **A, E ,F** Height parameters. **B, G** Number of unfurled leaves. **C, H** Leaf developmental plastochron, i.e., the leaf number that has just initiated from the SAM and is at the P1 stage. **D, I** the proportion of plants in which the SAM has differentiated into a floral meristem. **J** Fresh weight.

Bars represent mean ±SE, N=10. Asterisks indicate statistical significance from Mock treatment in an unpaired two-tailed t-test. **p<0.01, ***p<0.001, ****p<0.0001.

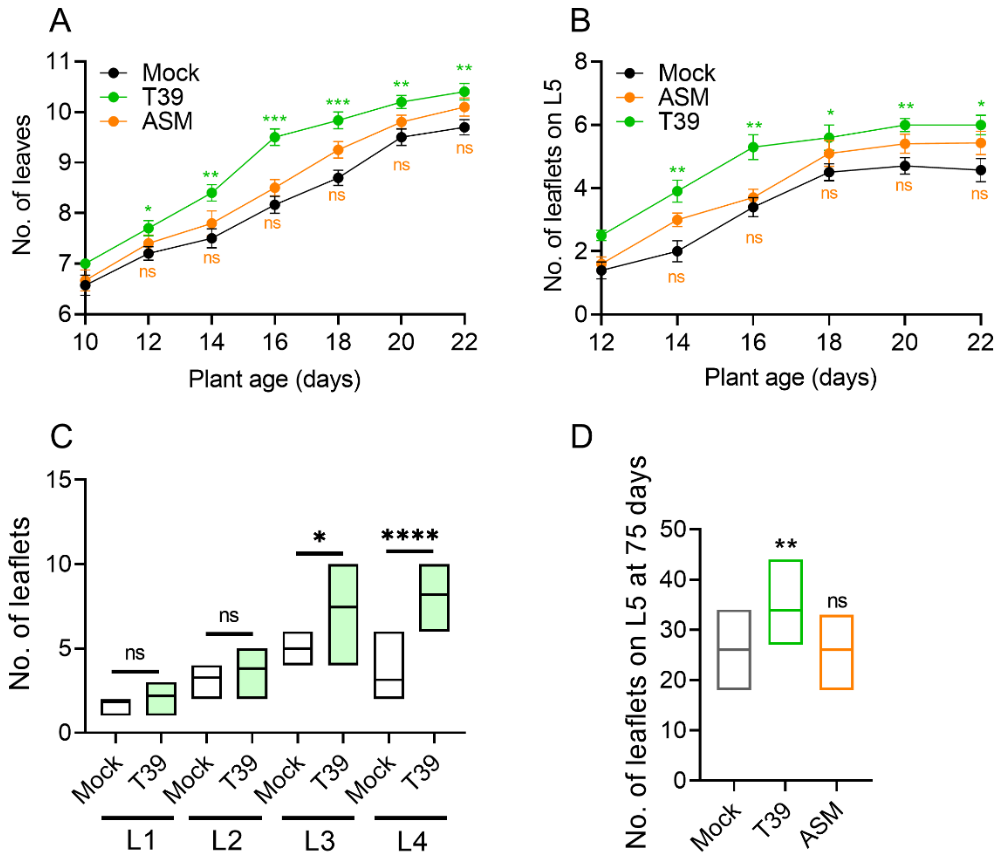


**Figure S4: Treatment with T39 accelerates leaf development.**

*S. lycopersicum* cv M82 Tomato seedlings were sprayed with elicitors twice (at 10 and 17 days of age, **A-C**), or soil-drenched 4 times, once a week, starting at the age of 3 weeks (**D**). Plants treated with DDW were used as mock. **A** Number of initiated leaves over time, starting from the day of the first elicitor treatment. **B** Number of leaflets on L5 over time, starting two days after the first elicitor treatment. **C** Number of leaflets on leaves 1-4, one week after the second treatment. **D** Number of leaflets on L5 of mature 75 day old plants.

**A-B** Points on kinetic graph represent mean ±SE, N=10. Asterisks indicate statistical significance from Mock treatment in multiple *t*-tests with Holm-Sidak correction. *p<0.05, **p<0.01, ***p<0.001, ns- non significant.

**C-D** Floating bars represent minimum to maximum values, line in box indicates mean, C: N=10, D: N=7. Asterisks indicate statistical significance from Mock treatment in unpaired two-tailed *t*-test. *p<0.05, **p<0.01, ****p<0.0001, ns- non significant.

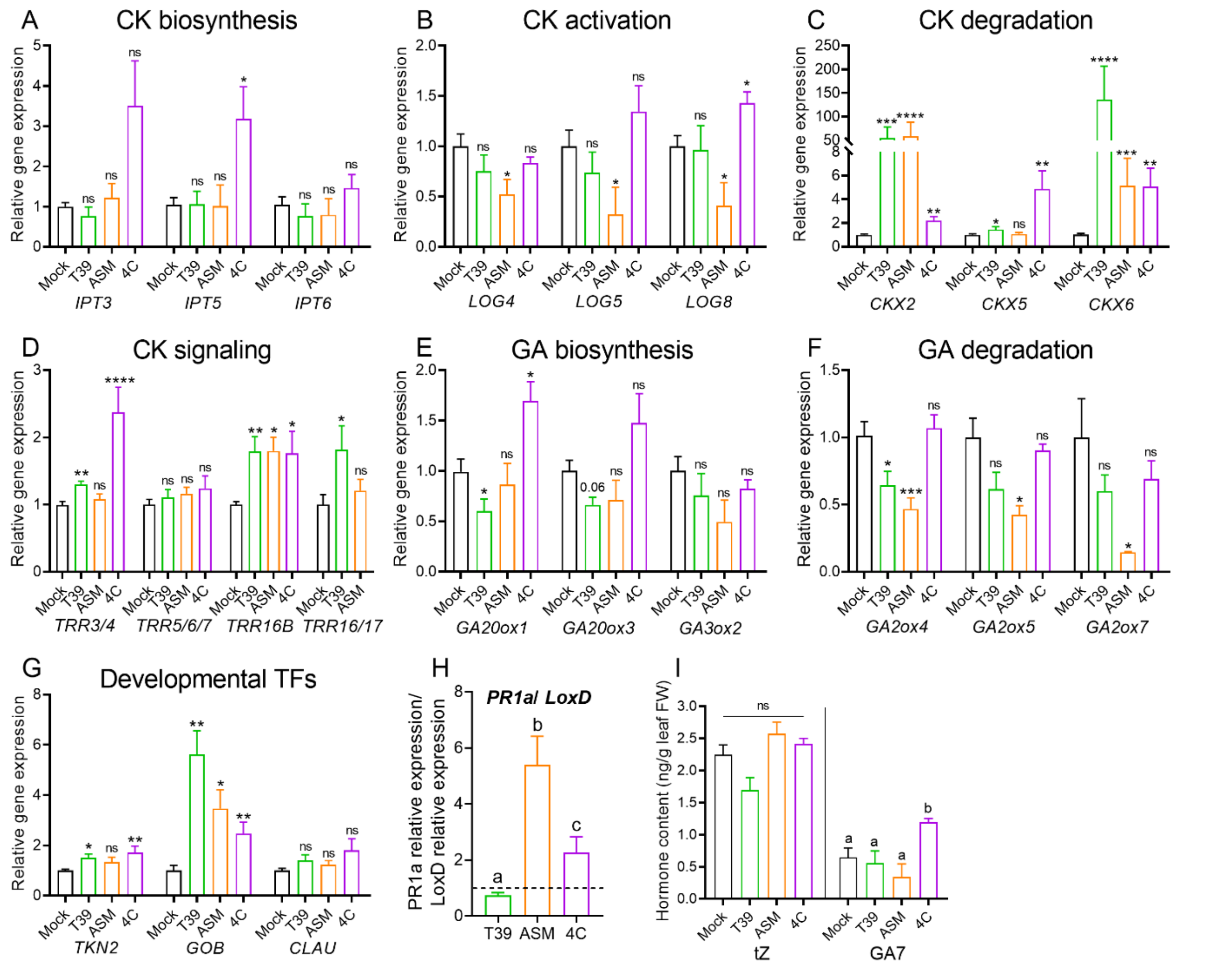


**Figure S5: Treatment with elicitors affects CK and GA pathway and developmental gene expression in sink leaves.**

*S. lycopersicum* cv Tomato seedlings were sprayed with elicitors twice, at 10 and 17 days of age. RNA was prepared from shhot apices (m+4), 48 h after the second treatment. Seedlings treated with DDW were used as mock. qRT-PCR was conducted to examine gene expression, with relative expression normalized to the geometric mean of the expression of 3 normalizer genes: *EXP (*Solyc07g025390*)*, *CYP* (Solyc01g111170), and *RPL8* (Solyc10g006580). **A** Cytokinin biosynthesis *IPT* genes. **B** Cytokinin activation *LOG* genes. **C** Cytokinin degradation *CKX* genes. **D** Cytokinin signaling response regulators (*TRRs*). **E** Gibberellin biosynthesis genes (*GA20ox* and *GA3ox*). **F** Gibberellin degradation genes (*GA2ox*). **G** Developmental transcription factors (*TKN2*- meristem maintenance; *GOB*- organ determination; *CLAU*- differentiation promotion). **H** Ratio of the expression of the defense genes *PR1a* and *LoxD*. A ratio below 1 suggests that ISR is the primary pathway activated, while a ratio above 1 suggests SAR activation. **I** Quantification of CK and GA derivatives in source leaf tissues. tZ- transZeatin, GA7- Gibberellin7.

Bars represent mean ±SE. A-G, I: N=8. A-G: Asterisks represent statistical significance from Mock treatment in a two-tailed *t*-test with Welch's correction where appropriate (unequal variances), for each gene. *p<0.05, **p<0.01, ***p<0.001, ****p<0.0001, ns- non significant. I: N=4. H-I: Different letters represent statistically significant differences among samples in Welch's ANOVA with Dunnett's post-hoc test (H), or in a two-tailed *t*-test with Welch's correction (I) where appropriate (unequal variances). H: p<0.042, I: p<0.018.

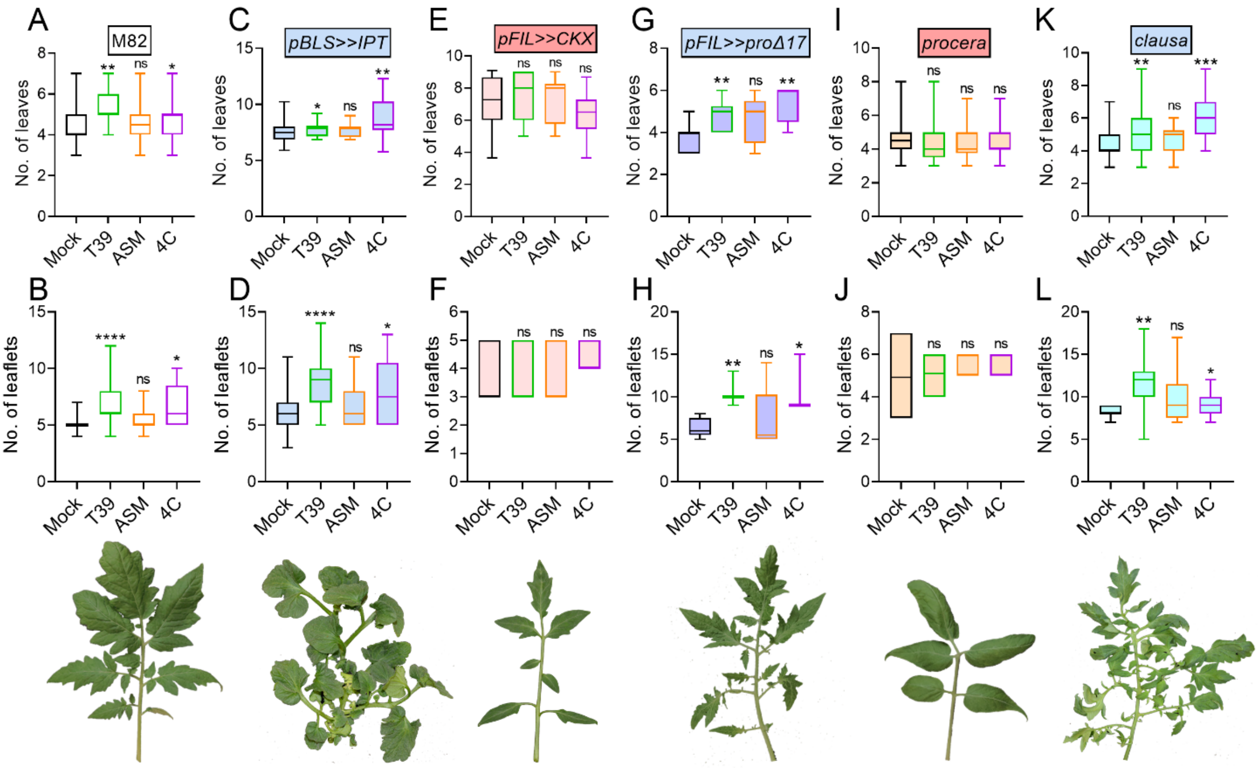


**<u>Figure S6: Treatment with T39 and 4C requires CK to promote leaf development.</u>**

Tomato seedlings of *S. lycopersicum* cv M82 (A-B), the transgenic *pBLS>>IPT* (C-D), *pFIL>>CKX* (E-F), *pFIL>>proΔ17* (G-H), or the recessive mutants *procera* (I-J) or *clausa* (K-L), all in the M82 background, were sprayed with elicitors twice, at 10 and 17 days of age. Seedlings treated with DDW were used as mock. Genotypes with high CK or low GA are indicated in blue, genotypes with low CK or high GA are indicated in red. High/ low refers to either content or signaling. See also <u>Table 1</u>. Representative images of a mature fifth leaf are provided below the graphs.

**A, C, E, G, I, K** No. of leaves, and **B, D, F, H, J, L**, No. of leaflets on L5, were counted 2 weeks after the second treatment.

Boxplots are shown with inter-quartile-ranges (box), medians (line in box) and outer quartile whiskers, minimum to maximum values. Asterisks indicate statistical significance from Mock treatment in one-way ANOVA with Bonferroni's post hoc test (A, D, F, K), Welch's ANOVA with Dunnett's post hoc test (B), or Welch's t-test (C, E, G, H, I, J, L). A: N=17. B: N=28. C: N=28. D: N=8. E: N=15. F: N=5. G: N=5. H: N=4. I: N=30. J: N=18. K: N=24. L: N=12. *$p<0.05$, **$p<0.01$, ***$p<0.001$, ****$p<0.0001$, ns- not significant.

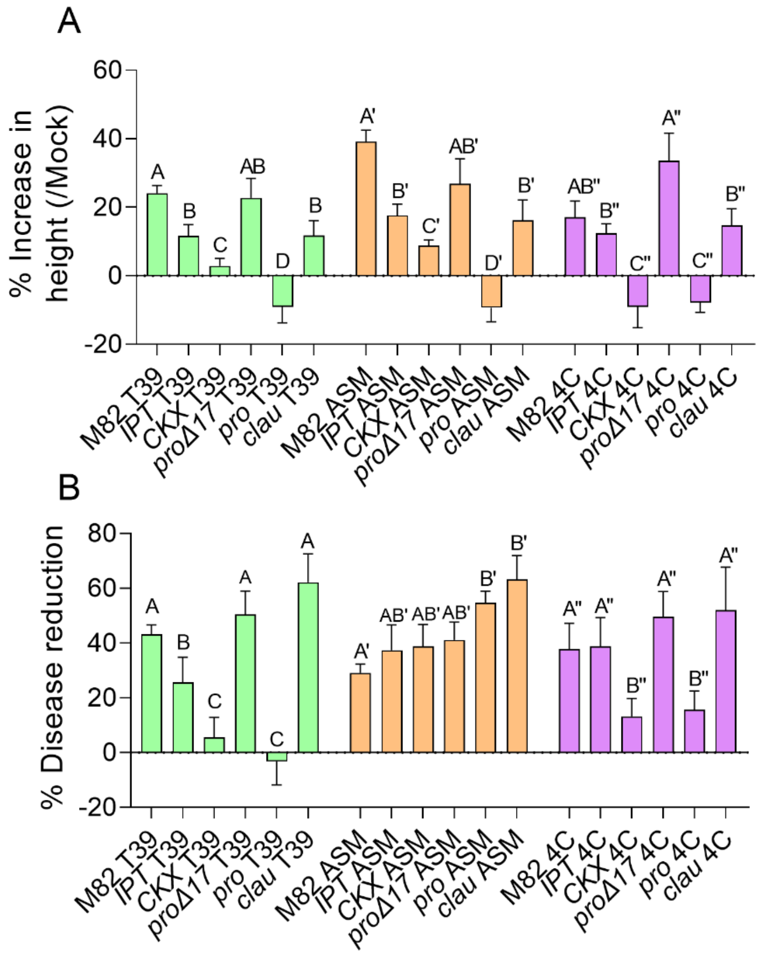


**Figure S7: Quantification of differential effects of elicitors on growth and disease resistance in altered CK and GA genotypes.**

Tomato seedlings of *S. lycopersicum* cv M82 (A-B), the transgenic *pBLS>>IPT*, *pFIL>>CKX*, *pFIL>>proΔ17*, or the recessive mutants *procera* or *clausa*, all in the M82 background, were sprayed with elicitors twice, at 10 and 17 days of age. Seedlings treated with DDW were used as mock.

**A** Height was measured 1 week after the second treatment.

**B** 4 week old tomato plants of the indicated genotypes were soil-drenched twice with elicitors: 3 days and 4 h before *Botrytis cinerea* inoculation. Plant were inoculated with 3 day-old *B. cinerea* mycelia. Plants treated with DDW were used as mock. Lesion area was measured 3 days after inoculation. Experiment was repeated 3 independent times.

Bars represent mean ±SE. Different letters indicate statistically significant differences between samples in a two-tailed *t*-test with Welch's correction: untagged letters for T39, tagged letters for ASM, and double-tagged letters for 4C. A: N≥5, p<0.048. B: N≥9, p<0.043.

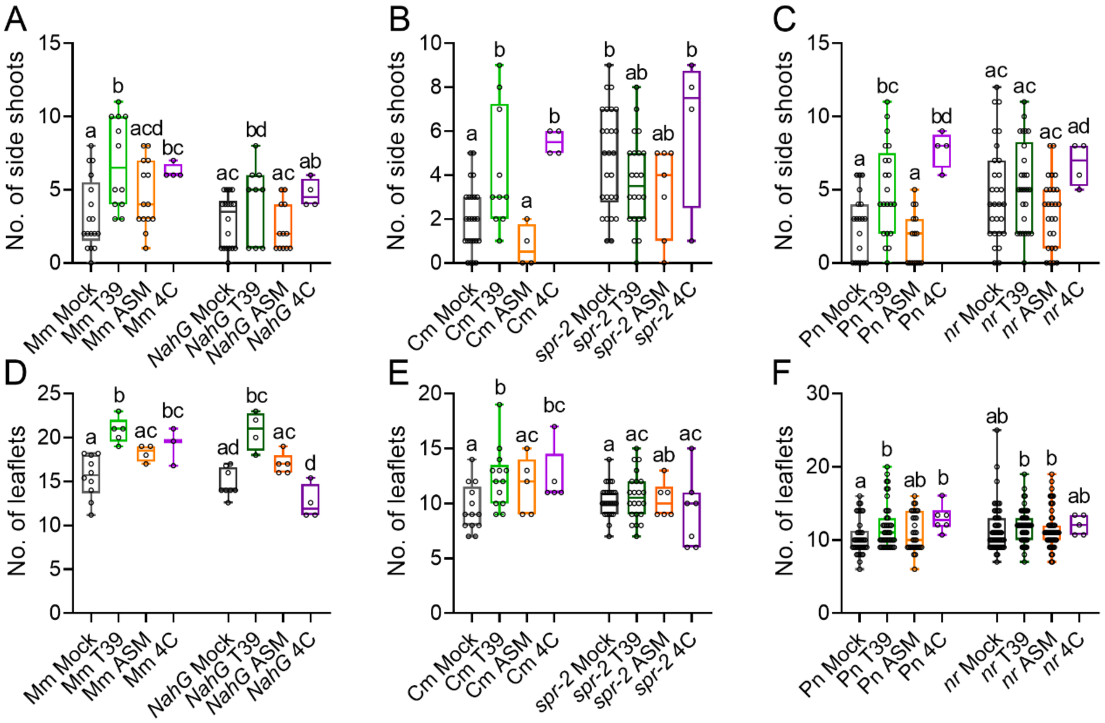


**Figure S8: Treatment with T39, ASM, and 4C has differential effects on development in defense pathway mutants.**

*S. lycopersicum* plants of the cultivars Moneymaker (MM) and the reduced SA transgene NahG (**A, D**), Castelmart (Cm) and the reduced JA mutant *spr-2* (**B, E**), and Pearson (Pn) and the reduced ET sensitivity mutant *neverripe* (*nr*) (**C, F**), were soil-drenched with elicitors once a week for four weeks.

**A-C** Side shoot production in 50 day old plants. **D-F** Leaf complexity of leaf No. 5 on 50 day old plants.

Box plots indicate inner quartile ranges (box), outer quartile ranges (whiskers), median (line), all points shown. Different letters indicate statistically significant differences among samples, in a two-tailed *t*-test with Welch's correction. A: N>4, p<0.0018. B: N>5, p<0.042. C: N>6, p<0.008. D: N>5, p<0.042. E: N>5, p<0.04. F: N>6, p<0.036.

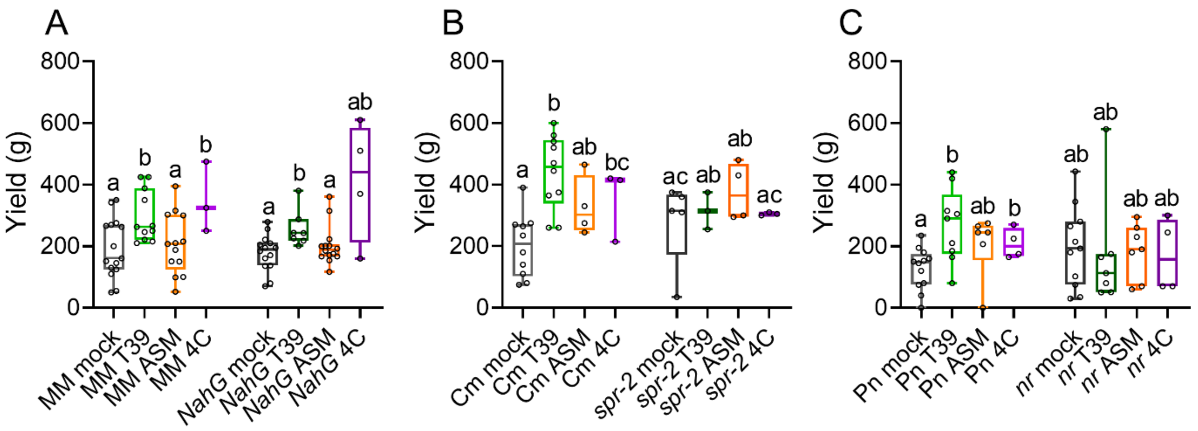


**<u>Figure S9: Treatment with T39 and ASM shows differential effects on yield in defense pathway mutants.</u>**

*S. lycopersicum* plants of the cultivars Moneymaker (MM) and the reduced SA transgene NahG (**A**), Castelmart (Cm) and the reduced JA mutant *spr-2* (**B**), and Pearson (Pn) and the reduced ET sensitivity mutant *neveripe* (*nr*) (**C**) were soil-drenched with elicitors once a week for four weeks. Plants treated with DDW were used as mock. Graphs depict average fruit weight per plant.

Boxplots are shown with inter-quartile-ranges (box), medians (line in box) and outer quartile whiskers, minimum to maximum values, all points shown. Different letters indicate statistically significant differences among samples, in a two-tailed *t*-test with Welch's correction, A: N>4, p< 0.046. B: N>3, p<0.036. C: N>4, p<0.045.

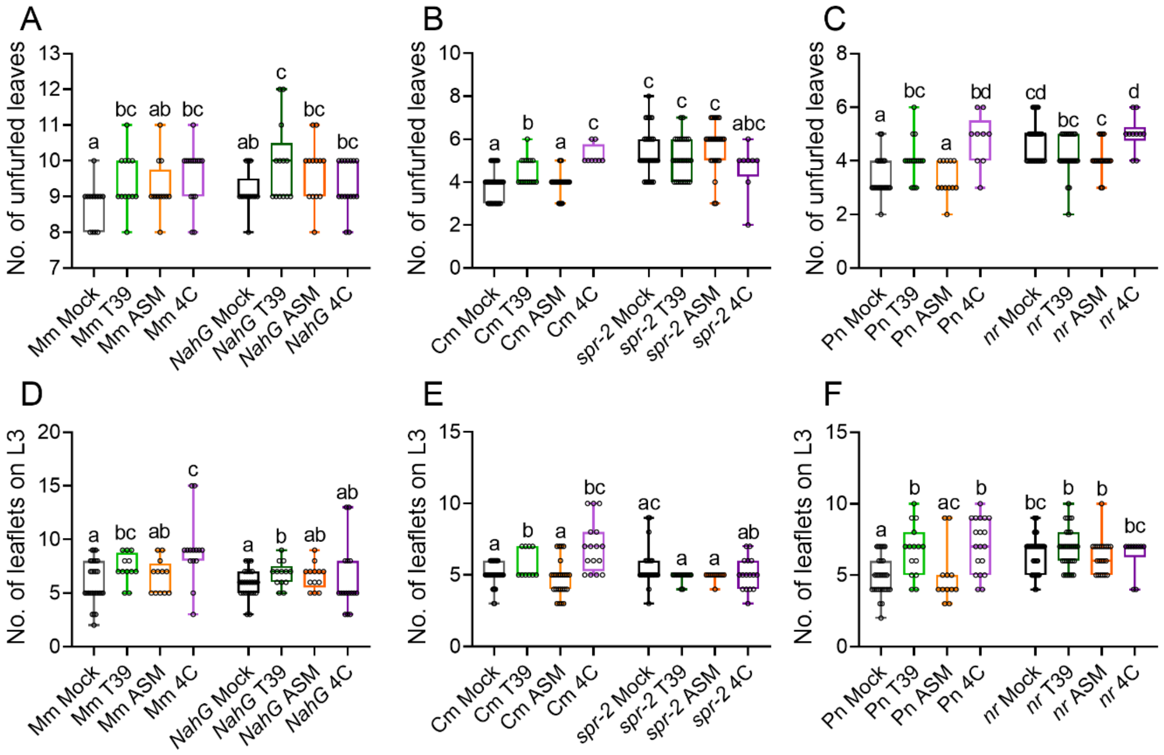


**Figure S10: Treatment with T39, ASM, or 4C shows differential effects on leaf development parameters in defense pathway mutant seedlings.**

*S. lycopersicum* seedlings of the cultivars Moneymaker (Mm) and the reduced SA transgene NahG (**A, D**), Castelmart (Cm) and the reduced JA mutant *spr-2* (**B, E**), and Pearson (Pn) and the reduced ET sensitivity mutant *neveripe* (*nr*) (**C, F**), were sprayed with elicitors twice, at 10 and 17 days of age. Seedlings treated with DDW were used as mock. Parameters were measured 1 week after the second treatment. **A-C** Number of leaves. **D-F** Leaf complexity, expressed as the number of leaflets on L3.

Boxplots are shown with inter-quartile-ranges (box), medians (line in box) and outer quartile whiskers, minimum to maximum values, all points shown, N>12. Different letters indicate statistically significant differences among samples in one-way ANOVA with Tukey's post hoc test, or in a two-tailed *t*-test.

A: N>12, p<0.036. B: N>8, p<0.028. C: N>9, p<0.029. D: N>12, p<0.028. E: N>10, p<0.027. F: N>12, p<0.042.

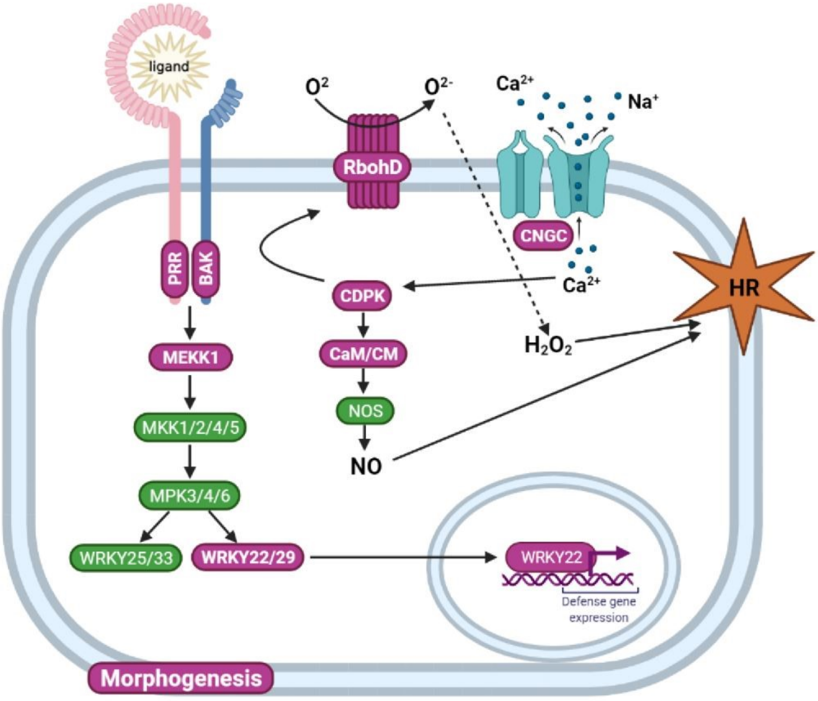


**Figure S11: Cartoon model of the overlap between developmental morphogenesis and defense responses.**

**The model details different genetic groups involved in defense responses within the cell**:

PRR- Pattern Recognition Receptor: recognition of MAMPs, PAMPs

Ligand- the PRR ligand could be a bacterial derived elicitor such as flagellin (**Fig. 1E**), or a fungal-derived elicitor such as EIX (**Fig. 1D**).

BAK- Bri-Associated Kinase: immune co-receptor for several PRRs

MEKK/ MKK/ MPK- Mitogen activated protein kinases: transduction of immune signals

WRKY- Transcriptional regulation of biotic stress responses

RbohD- Membrane localized NADPH oxidase: generation of ROS

CDPK- Calcium Dependent protein Kinase: stress response

CNGC- cyclic nucleotide-gated channel: PAMP-induced calcium signaling

CaM/ CM- Calmodulin: Calcium sensors, involved in calcium-dependent regulation of gene expression during plant immune responses.

NOS- Nitric Oxide Synthase: required for HR, involved in both SA and ET/JA signaling

**Generated defense responses**: Defense gene expression (see **Fig. 3**), Reactive oxygen species (such as $H_2O_2$) (see **Fig. 1**), Nitric Oxide (NO), Hypersensitive response (HR).

**Gene groups highlighted in fuchsia are also found in the group of genes promoting morphogenesis** (Israeli et al. 2021). Illustration created with BioRender.com.

**Supplemental Table 1: Genes recognized as "plant-pathogen interaction" pathway genes by KEGG, that are defined as morphogenetic.**

Listed are genes belonging to the morphogenetic dataset (Israeli et al., 2021) that were recognized as belonging to the KEGG group "plant-pathogen interaction" (Fishers exact test, $p<0.05$). The table lists the relative expression levels of each gene in successive developmental stages.

| Gene_ID | Description | m+P3 | P4-d | P5-d | P6-d |
|---|---|---|---|---|---|
| Solyc10g074570 | Calcium dependent protein kinase 6 (AHRD V1 **** B9MZ11_POPTR); contains Interpro domain(s) IPR002290 Serine/threonine protein kinase | 79.0 | 50.7 | 40.0 | 29.3 |
| Solyc04g015980 | FLS2 homolog Receptor kinase-like protein (AHRD V1 **** Q7DMC2_ORYLO); contains Interpro domain(s) IPR002290 Serine/threonine protein kinase | 11.3 | 7.0 | 5.3 | 4.3 |
| Solyc12g010010 | Cyclic nucleotide gated channel (AHRD V1 ***- A9CRE4_MALDO); contains Interpro domain(s) IPR000595 Cyclic nucleotide-binding | 125.7 | 117.7 | 102.0 | 89.3 |
| Solyc03g083520 | Calmodulin (AHRD V1 ***- B6T4U8_MAIZE); contains Interpro domain(s) IPR011992 EF-Hand type | 68.0 | 63.7 | 58.7 | 52.3 |
| Solyc07g065840 | Heat shock protein 90 -2 | 8570.3 | 7991.7 | 5098.0 | 4105.7 |
| Solyc12g099990 | SlCaM5 is a member of the calmodulin gene family which modulates the response to calcium signalling. Calmodulin 2 (AHRD V1 ***- Q710C9_BRAOL); contains Interpro domain(s) IPR011992 EF-Hand type | 417.0 | 315.7 | 260.3 | 222.3 |
| Solyc11g071740 | Calmodulin-like protein | 19.7 | 7.7 | 5.3 | 1.3 |
| Solyc04g018110 | Calmodulin-like protein 1 | 43.0 | 38.3 | 21.7 | 19.0 |
| Solyc01g008950 | CaM1 is a member of the Calmodulin gene family which modulate responses to calcium signalling. Calmodulin 5/6/7/8-like protein | 328.7 | 277.7 | 219.7 | 213.3 |
| Solyc01g104970 | BAK/SERK3B Receptor-like kinase (AHRD V1 **** A7VM44_MARPO); contains Interpro domain(s) IPR002290 Serine/threonine | 49.3 | 48.3 | 46.7 | 46.3 |

| | | | | | |
|---|---|---|---|---|---|
| | protein kinase Receptor like kinase, RLK | | | | |
| Solyc03g098050 | SlCaM6 is a member of the calmodulin gene family which modulates the response to calcium signalling. Calmodulin 3 protein (AHRD V1 **** Q712P2_CAPAN); contains Interpro domain(s) IPR011992 EF-Hand type | 297.0 | 234.0 | 169.3 | 169.0 |
| Solyc04g009800 | Calcium-dependent protein kinase 2 (AHRD V1 **** Q93YF4_TOBAC); contains Interpro domain(s) IPR002290 Serine/threonine protein kinase | 72.0 | 71.3 | 62.0 | 60.0 |
| Solyc08g081690 | RBOH1 NADPH oxidase | 13.7 | 5.0 | 4.7 | 1.7 |
| Solyc07g053170 | SlMAPKKK56 is a part of the MAPKKK gene family known to be involved in the response to various biotic and abiotic stresses in plants. Protein serine/threonine kinase | 109.3 | 105.0 | 85.7 | 82.7 |
| Solyc12g098820 | Pto homolog Receptor-like kinase (AHRD V1 ***- A7VM24_MARPO); contains Interpro domain(s) IPR002290 Serine/threonine protein kinase | 127.7 | 126.0 | 121.7 | 115.0 |
| Solyc01g104530 | SlMAPKKK10 is a part of the MAPKKK gene family known to be involved in the response to various biotic and abiotic stresses in plants. Protein serine/threonine kinase | 103.0 | 100.3 | 87.3 | 82.7 |
| Solyc01g095100 | SlWRKY22 is a member of WRKY transcription factor gene family which has been implicated in multiple biological processes in plants. It has the C2H2 Zinc-finger domain. Computational annotation: WRKY transcription factor 23 (AHRD V1 ***- C9DI12_9ROSI); contains Interpro domain(s) IPR003657 DNA-binding WRKY | 14.0 | 10.0 | 8.0 | 7.0 |

**Supplemental Table 2: qPCR primers used in this work**

| Locus | Name | Forward | Reverse |
|---|---|---|---|
| Solyc07g025390 | *EXP* | TGGGTGTGCCTTTCTGAATG | GCTAAGAACGCTGGACCTAATG |
| Solyc10g006580 | *RPL8* | TGGAGGGCGTACTGAGAAAC | TCATAGCAACACCACGAACC |
| Solyc01g111170 | *CYP* | TGAGTGGCTCAACGGAAAGC | CCAACAGCCTCTGCCTTCTTA |
| Solyc01g106620 | *PR1a* | CTGGTGCTGTGAAGATGTGG | TGACCCTAGCACAACCAAGA |
| Solyc03g122340 | *LoxD* | CCATCCTCACCACCCTCATC | TACTCGGGATCGTTCTCGTC |
| Solyc01g088160 | *CKX2* | CCCCGAAAATGGTGAAATG | CAAAGTGGCTTGCTTGAACA |
| Solyc04g016430 | *CKX5* | TGTCACTGGTAAAGGAGAGGTG | GAGCAATCCTAGCCCTTGTG |
| Solyc12g008900 | *CKX6* | CAGGTGCTAAGCCATACTCTAGG | GGACATTCCATTAGGGGACA |
| Solyc01g080150 | *IPT3* | TTCCATGCTTGATGTGCTTC | GCTTGCTGTCAACGTCAAAA |
| Solyc11g066960 | *IPT5* | CCAGTTGCAGCCAACAGAAT | GACCCCAGTTCTGCCCTCTA |
| Solyc12g014190 | *IPT6* | GATGTTCCAAAAGCCTCTCG | TAAACTTGCAAGCTCTGAGTCG |
| Solyc04g081290 | *LOG3* | TGGGCCTAGTTTCTCAATCAG | TTTAGGAATCACCCCTAACACG |
| Solyc01g005680 | *LOG4* | GACAAAGGTGTGGAAGAAGGA | GCTCTTTGGGTGATGTAGCTG |
| Solyc08g062820 | *LOG5* | CATGTTGCTCCCCATGAAA | TGGAGACTGCTCCTTTGGAT |
| Solyc06g075090 | *LOG8* | TGAGCTTGGAAGAGAAATAGTATCAA | AAACCCATCAAACCAATGCT |
| Solyc05g006420 | *TRR3/4* | CGTCCCCTAAAGCATTCTCA | CGTCTTGTTGGTGATGTTGG |
| Solyc03g113720 | *TRR5/6/7* | GGGATTGATGGTTTGAAGGT | ATCTTGCTCAACACCGATGA |
| Solyc06g048600 | *TRR 16b* | CATCAATGCATGGAAGAAGG | GCATTGCATTATTTGGCATC |
| Solyc03g006880 | *GA20ox1* | AGATTGTGTTGGTGGACTTCAA | TAGCGCCATAAATGTGTCG |
| Solyc11g072310 | *GA20ox3* | ACTTTAGGGACAGGGCCTCA | ACTTGAAGCCCACCAACACT |
| Solyc03g119910 | *GA3ox2* | TTGGCCATGCATGCAAAACA | ATCTCGTCCCGTGTGTTTCC |
| Solyc07g061720 | *GA2ox4* | CCAACAACACTTCCGGTCTT | CATTCGTCATCACCTGTAATGAG |
| Solyc07g061730 | *GA2ox5* | CAACACATCCGGCCTTCAAA | AACTCTGATCAGGTGGCACA |
| Solyc02g080120 | *GA2ox7* | AGCCACCTCCACTTCTCAAT | GGTTTGGCTGCTGTGACAAG |
| Solyc02g081120 | *TKN2* | CCATATCCATCGGAATCTCAG | TGGTTTCCAATGCCTCTTTC |
| Solyc07g062840 | *GOB* | CAGGAGTTCGAAGGACGAGTGG | TTGGCTGTAGTGTATGCAAGGTG |
| Solyc04g008480 | *CLAU* | CCTCTCACAACAAGCAATGAACTT | AGGACGATGCAATGAGAGAGAC |

**Supplemental Table 3: Genes assayed in qPCR and their expected expression**

| Locus | Name | Gene group | Expected expression** | Comments |
|---|---|---|---|---|
| Solyc07g025390 | *EXP* | Normalizer | Strong, ubiquitous | |
| Solyc10g006580 | *RPL8* | Normalizer | Strong, ubiquitous | |
| Solyc01g111170 | *CYP* | Normalizer | Strong, ubiquitous | |
| Solyc01g106620 | *PR1a* | Defense (SAR) | Increased following elicitation | Detected in both sink and source leaf tissues |
| Solyc03g122340 | *LoxD* | Defense (ISR) | Increased following elicitation | Detected in both sink and source leaf tissues |
| Solyc08g061930 | *CKX1* | CK degradation | Low expression in young and mature leaves | Not detected in qPCR |
| Solyc01g088160 | *CKX2* | CK degradation | Low expression in young and mature leaves | Detected in both sink and source leaf tissues |
| Solyc08g061920 | *CKX3* | CK degradation | Low expression in young leaves, zero expression in mature leaves | Not detected in qPCR |
| Solyc04g080820 | *CKX4* | CK degradation | Undetected in young leaves, zero expression in mature leaves | Not detected in qPCR |
| Solyc04g016430 | *CKX5* | CK degradation | Undetected in young leaves, low expression in mature leaves | Detected in both sink and source leaf tissues |
| Solyc12g008900 | *CKX6* | CK degradation | Zero expression in both young and mature leaves | Detected only in sink leaf tissues |
| Solyc10g079870 | *CKX7* | CK degradation | Undetected in young leaves, zero expression in mature leaves | Not detected in qPCR |
| Solyc05g009410 | *IPT1* | CK biosynthesis | Very low expression in young and mature leaves | Not detected in qPCR |
| Solyc04g007240 | *IPT2* | CK biosynthesis | Very low expression in young leaves, zero expression in mature leaves | Not detected in qPCR |
| Solyc01g080150 | *IPT3* | CK biosynthesis | Low expression in young and mature leaves | Detected in both sink and source leaf tissues |
| Solyc09g064910 | *IPT4* | CK biosynthesis | Zero expression in both young and mature leaves | Not detected in qPCR |
| Solyc11g066960 | *IPT5* | CK biosynthesis | Low expression in young and mature leaves | Detected in both sink and source leaf tissues |
| Solyc12g014190 | *IPT6* | CK biosynthesis | Undetected in young leaves, low expression in mature leaves | Detected in both sink and source leaf tissues |
| Solyc11g069570 | *LOG1* | CK activation | Low expression in young leaves, strong | Unable to obtain or generate satisfactory primers |

| | | | | |
|---|---|---|---|---|
| | | | expression in mature leaves | |
| Solyc09g007830 | *LOG2* | CK activation | Expressed in young and mature leaves | Unable to obtain or generate satisfactory primers |
| Solyc04g081290 | *LOG3* | CK activation | Very low expression in young leaves, expressed in mature leaves | Detected only in source leaf tissues |
| Solyc01g005680 | *LOG4* | CK activation | Very low expression in young leaves, expressed in mature leaves | Detected in both sink and source leaf tissues |
| Solyc08g062820 | *LOG5* | CK activation | Low expression in young and mature leaves | Detected in both sink and source leaf tissues |
| Solyc10g084150 | *LOG6* | CK activation | Low expression in young leaves, expressed in mature leaves | Unable to obtain or generate satisfactory primers |
| Solyc10g082020 | *LOG7* | CK activation | Very low expression in young leaves, zero expression in mature leaves | Not detected in qPCR |
| Solyc06g075090 | *LOG8* | CK activation | Low expression in young leaves, expressed in mature leaves | Detected in both sink and source leaf tissues |
| Solyc05g006420 | *TRR3/4* | CK response | Expressed in young leaves, low expression in mature leaves | Detected only in sink leaf tissues |
| Solyc03g113720 | *TRR5/6/7* | CK response | Expressed in young and mature leaves | Detected in both sink and source leaf tissues |
| Solyc10g079600 | *TRR8/9a* | CK response | Undetected in young leaves, low expression in mature leaves | Not tested |
| Solyc02g071220 | *TRR8/9b* | CK response | Expressed in young and mature leaves | Unable to obtain or generate satisfactory primers |
| Solyc10g079700 | *TRR8/9c* | CK response | Undetected in young leaves, low expression in mature leaves | Not tested |
| Solyc06g048930 | *TRR 16/17* | CK response | Very low expression in young and mature leaves | Detected only in sink leaf tissues |
| Solyc06g048600 | *TRR 16b* | CK response | Expressed in young leaves, low expression in mature leaves | Detected in both sink and source leaf tissues |
| Solyc03g006880 | *GA20ox1* | GA biosynthesis | Very low expression in young leaves, expressed in mature leaves | Detected in both sink and source leaf tissues |
| Solyc06g035530 | *GA20ox2* | GA biosynthesis | Very low expression in young and mature leaves | Not detected in qPCR |
| Solyc11g072310 | *GA20ox3* | GA biosynthesis | Very low expression in young and mature leaves | Detected in both sink and source leaf tissues |
| Solyc03g119910 | *GA3ox2* | GA biosynthesis | Very low expression in young and mature leaves | Detected in both sink and source leaf tissues |

| | | | | |
|---|---|---|---|---|
| Solyc07g061720 | *GA2ox4* | GA degradation | Very low expression in young and mature leaves | Detected in both sink and source leaf tissues |
| Solyc07g061730 | *GA2ox5* | GA degradation | Very low expression in young and mature leaves | Detected in both sink and source leaf tissues |
| Solyc02g080120 | *GA2ox7* | GA degradation | Low expression in young leaves, expressed in mature leaves | Detected in both sink and source leaf tissues |
| Solyc02g081120 | *TKN2* | Transcription factor | Very low expression in young and mature leaves | Detected only in sink leaf tissues |
| Solyc07g062840 | *GOB* | Transcription factor | Expressed in young leaves, zero expression in mature leaves | Detected only in sink leaf tissues |
| Solyc04g008480 | *CLAU* | Transcription factor | Low expression in young leaves, zero expression in mature leaves | Detected only in sink leaf tissues |

Genes indicated in gray were not included in the manuscript.

** Putative expression provided by:

(1) Martinez CC, Li S, Woodhouse MR, Sugimoto K, Sinha NR. Spatial transcriptional signatures define margin morphogenesis along the proximal–distal and medio-lateral axes in tomato (*Solanum lycopersicum*) leaves. *The Plant Cell*, 2021, https://doi.org/10.1093/plcell/koaa012.

(2) Fernandez-Pozo N, Zheng Y, Snyder SI, Nicolas P, Shinozaki Y, Fei Z, Catala C, Giovannoni JJ, Rose JKC, Mueller LA. The Tomato Expression Atlas. *Bioinformatics,* 2017, doi: 10.1093/bioinformatics/btx190.
